# Cryo-EM structures capturing the entire transport cycle of the P4-ATPase flippase

**DOI:** 10.1101/666321

**Authors:** Masahiro Hiraizumi, Keitaro Yamashita, Tomohiro Nishizawa, Osamu Nureki

## Abstract

In eukaryotic membranes, P4-ATPases mediate the translocation of phospholipids from the outer to inner leaflet and maintain lipid asymmetry, which is critical for protein trafficking and signaling pathways. Here we report the cryo-EM structures of six distinct intermediates of the human ATP8A1-CDC50a hetero-complex, at 2.6–3.3 Å resolutions, revealing the entire lipid translocation cycle of this P4-ATPase. ATP-dependent phosphorylation induces a large rotational movement of the actuator domain around the phosphorylation site, accompanied by lateral shifts of the first and second transmembrane helices, thereby allowing phosphatidylserine binding. The phospholipid head group passes through the hydrophilic cleft, while the acyl chain is exposed toward the lipid environment. These findings advance our understanding of the flippase mechanism and the disease-associated mutants of P4-ATPases.

**One Sentence Summary:** Cryo-EM reveals lipid translocation by P4-type flippase.

## Main Text

### Introduction

In eukaryotic cells, the phospholipid compositions differ between the outer and inner leaflets of the plasma/organellar membranes: phosphatidylcholine (PC) and sphingomyelin are enriched in the outer leaflet, whereas phosphatidylserine (PS) and phosphatidylethanolamine (PE) are confined to the inner leaflet. The maintenance and disruption of the asymmetric composition affect fundamental processes, such as membrane biogenesis, membrane trafficking, signaling, and apoptosis. Three types of transporters, scramblase, floppase, and flippase, have been suggested to function as phospholipid translocators. While scramblase catalyzes the energy independent and bidirectional phospholipid translocation that dissipates membrane asymmetry, flippase and floppase mediate specific phospholipid translocations against their concentration gradients, utilizing the energy of ATP hydrolysis to maintain the asymmetric phospholipid composition. ATP-binding cassette (ABC) transporters function as floppases that drive the inner-to-outer translocation of lipids, whereas Type IV P-type ATPases (P4-ATPase) are flippases that drive the outer-to-inner translocation of lipids (*1*–*4*).

The transport by the P-type ATPases occurs essentially according to the Post-Albers mechanism, wherein ATP hydrolysis-coupled phosphorylation and dephosphorylation within the cytoplasmic ATPase domain mediate the transition between the two intermediate states, E1 and E2, which have different affinities for the substrates, enabling the substrate transport across the membrane (*5, 6*). Among the P-type ATPase family members, the P1- to P3-ATPases are ion transporters, with the most representative being the P2-ATPase family, including the sarcoplasmic reticulum Ca^2+^ pump (SERCA), Na^+^/K^+^-ATPase, and H^+^/K^+^-ATPase, whereas the P4-ATPase is the only member that functions as a lipid transporter (*7*). The human genome encodes 14 P4-ATPase subclasses, which differ in their lipid selectivities and tissue expression (*8*). Most of the P4-ATPases form heterodimers with the CDC50 family protein, which is essential for the proper expression and flippase activity of the P4-ATPases (*9, 10*).

The first identified P4-ATPase member, ATP8A1, was found in bovine erythrocytes and chromaffin granules as an aminophospholipid translocase (*11, 12*). P4-ATPases, including ATP8A1, are present in plasma/organellar membranes and sequester phosphoserine (PS) lipids (known as the “eat-me-signal”) from the outer- to inner-leaflet to allow resting cells to escape from phagocytosis, whereas in apoptotic cells, their cleavage and inactivation by proteases, such as caspases and calpains, lead to the PS exposure on the cell surface and thus induce phagocytosis (*13, 14*). Furthermore, the ATP8A1-catalyzed flipping of PS in the organellar membrane is necessary for the transport of recycling endosomes, membrane fission, and cell migration (*15, 16*). Several diseases are associated with P4-ATPases. For example, ATP8B1 mutations cause the liver diseases known as BRIC1 and PFIC1, ATP10A is associated with type 2 diabetes and insulin resistance, and ATP11A is associated with cancer (*17*). Furthermore, ATP8A1 and ATP8A2 have been identified as causative genes for neurological disorders. ATP8A1 knockout mice show hippocampus-dependent learning deficits, associated with the exposure of PS on the outer surface of the plasma membrane in hippocampal neurons (*18*), and missense mutations (I376M and N917D) of ATP8A2 have also been identified as the cause of the rare neurodegenerative disease known as cerebellar ataxia, mental retardation, and dysequilibrium syndrome (CAMRQ) (*19, 20*).

As compared to the canonical ion-transporting P-type ATPases, P4-ATPase has a large transport substrate, and thus is expected to utilize a different mechanism for the substrate recognition and translocation (*21*–*23*). However, despite substantial efforts, the molecular mechanism underlying the lipid flippase activity by the P4-ATPases has remained elusive. Here we report the cryo-EM structures of the human ATP8A1-CDC50a heterodimer complex in the absence of ligand (E1) and in the presence of an ATP analogue (E1-ATP), AlF_4_^−^-ADP (E1P-ADP), BeF_3_^−^ (E2P), and AlF_4_^−^ (E1P and phospholipid-bound E2P), revealing the entire transport cycle along the lipid flipping reaction. These findings advance our understanding of the dynamic phospholipid translocation mechanism of P4-ATPase.

### Overall structure

We performed a cryo-EM analysis of the P4-ATPase lipid translocator family to elucidate the lipid translocation mechanism (Fig. 1). We expressed full-length human ATP8A1 and human CDC50a together in mammalian HEK293F cells, and the complex was purified in GDN micelles (fig. S1A). The SDS-PAGE analysis showed multiple bands of CDC50a, indicating the heterogeneous glycosylation (fig. S1B). The purified ATP8A1-CDC50a complex showed PS-dependent ATPase activity, with a *K*_m_ of 111 ± 26.4 µM and a *V*_max_ of 99.7 ± 9.50 nmol min^-1^mg^−1^, as well as weak PE-dependent ATPase activity (Fig. 1C), consistent with the previous reports (*12, 16*). The ATPase activity was inhibited by the general inhibitors of P-type ATPases, such as BeF_3_^−^ and AlF_4_^−^ (fig. S1D). The purified ATP8A1-CDC50a complex was subjected to cryo-EM single particle analyses under several different conditions; namely, without any inhibitors and in the presence of AMP-PCP, ALF_4_^−^-ADP, BeF_3_^−^, and ALF_4_^−^. The acquired movies were motion-corrected and processed in RELION 3.0 (*24*), which provided cryo-EM maps at overall resolutions of 2.6–3.3 Å, according to the gold-standard Fourier shell correlation (FSC) 0.143 criterion (figs. S4-8). The flexible cytoplasmic ATPase domain is most stabilized in the AlF_4_^−^-ADP and BeF_3_^−^-bound states, allowing the *de novo* modeling of almost the entire ATP8A1-CDC50a complex, except for some minor disordered regions (Fig. 1, A, B and fig. S9). The overall structure shows the typical P-type ATPase fold, composed of three large cytoplasmic domains (A, actuator; N, nucleotide binding; P, phosphorylation) and ten membrane-spanning helices (M1-10). CDC50a has two transmembrane helices (TM1 and TM2) at the N- and C-termini, an ectodomain consisting of an antiparallel β-sandwich (β1-8), and extensions of about 60 and 70 amino acids with less secondary structure in the β3-4 and β5-6 loops, respectively, which are stabilized by two intra-chain disulfide bonds (fig. S3C). The three N-linked glycosylation sites of CDC50a, which are important for the proper folding as well as the membrane trafficking of the P4-ATPases, are clearly visible in the cryo-EM map (fig. S3C, D) (*10, 25*).

**Figure 1:**
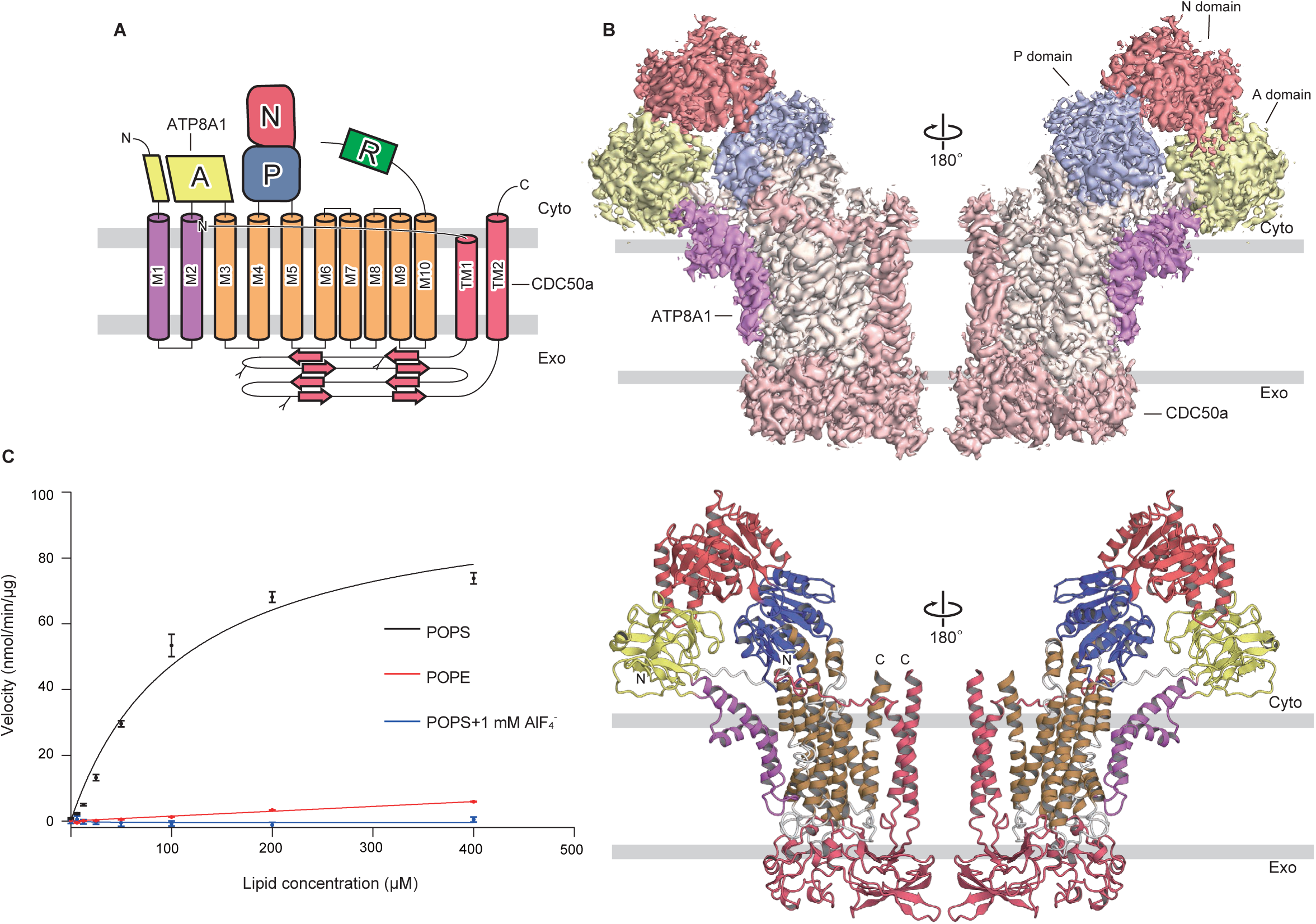
Biochemical and Cryo-EM studies of the ATP8A1-CDC50a complex. (**A**) Topology diagram of ATP8A1-CDC50a. Conserved domains and transmembrane helices are schematically illustrated. In the cytoplasmic regions, the A-, N-, and P-domains and the C-terminal regulatory domain are colored yellow, red, blue, and light green, respectively. M1-2 and M3-10 of ATP8A1 are purple and brown, respectively, while CDC50a is pink. The N-glycosylation sites are shown as sticks. cyto, cytoplasmic side; exo, exoplasmic side. (**B**) Overall structure of ATP8A1-CDC50a. Cryo-EM maps of the ATP8A1-CDC50a complex are shown on the top and the ribbon models are below. The same color scheme is used throughout the manuscript. (**C**) Phospholipids-dependent ATPase activities of ATP8A1. Data points represent the means ± SEM of three to six measurements at 37°C. By nonlinear regression of the Michaelis-Menten equation, ATP8A1-CDC50a in GDN micelles has a *K*_m_ of 111.0 ± 26.4 μM for POPS and a maximal ATPase activity of 99.7 ± 9.5 nmol/mg/min.

### Interaction between ATP8A1 and CDC50a

CDC50a is an essential component for the P4-ATPases and is required for the proper expression and folding of ATP8A1 (fig. S1C) (*9, 10*). CDC50a and ATP8A1 interact extensively through the extracellular transmembrane and intracellular regions (fig. S3). In the extracellular region, the CDC50a ectodomain covers all of the extracellular loops of ATP8A1, except for the M1-2 loop, interacting in an electrostatic complementary manner: the extracellular loops of ATP8A1 bear negative charges, whereas CDC50a bears positive charges (fig. S3A). In particular, Asp961 and Glu1026 of ATP8A1 form a salt bridge with Arg262 of CDC50a. In addition, the M3-4 loop of ATP8A1 extends toward CDC50a, and the two bulky residues at the tip of the loop, Trp328 and Tyr329, form hydrophobic interactions involving Phe127, Tyr299, Pro300, Val301, and the N-glycan attached to Asn180 of CDC50a (fig. S3D). In the transmembrane region, several bulky residues, such as Trp942, Ala947 (M9), Met1038, Phe1042, and Leu1049 (M10) of ATP8A1 and Phe54, Ile57, Phe61 (TM1) and Phe324, Leu325, Ala328, Tyr329, and Val332 (TM2) of CDC50, are engaged in the complex interaction (fig. S3F). Furthermore, we observed a strong planar density at the interface between M7 and M10 of ATP8A1 and TM2 of CDC50a (fig. S3B), which could be assigned to the cholesteryl hemisuccinate (CHS) added during solubilization. Therefore, cholesterol may bind to the same site and facilitate the hetero-dimeric interaction of ATP8A1 and CDC50a. In the cytoplasmic region, the N-terminal tail of CDC50a adopts an unstructured loop conformation that extends parallel to the plasma membrane and interacts with the M6-7 and M8-9 loops and the short segment connecting M4 and the P-domain (fig. S3E). Overall, CDC50a envelops the bulk of the TM segments and forms extensive interactions with ATP8A1, which explains the chaperone activity of CDC50a for the P4-type ATPases.

### Entire transport cycle of P4-ATPase

The cryo-EM structures showed the clear densities of the inhibitors in their respective maps, bound at the catalytic site of ATP8A1 and stabilizing the ATPase domain in different conformations (Fig. 2), whereas CDC50a adopted almost the same conformation in all of these states. The structures obtained under three conditions, namely without inhibitor, with AMP-PCP, and with AlF_4_^−^-ADP, describe the conformational changes upon ATP binding and auto-phosphorylation, which correspond to the E1, E1-ATP, and E1P-ADP conformations, respectively, in the Post-Albers scheme (Fig. 2A & Movie S1). The densities of the N- and A-domains are only weakly visible in the E1 state, indicating the highly flexible motion of these domains without any ligand (fig. S4). The particles were then classified according to the densities of the N- and P-domains, and these domains were modeled into the class with the strongest densities, which probably represents the most likely arrangement in the E1 state (class 2 in fig. S4). The particles of the AMP-PCP-bound state can be classified into three similar conformations, wherein the N- and P-domains adopt slightly different orientations (fig. S5). The comparison of these classes indicated that ATP binding at the N-domain induces the mutual approaching of the N- and P-domains. The density for the AMP-PCP is most clearly visible with the proximal arrangement of these domains, in which ATP bridges the two domains (fig. S5D): the adenine ring interacts with Phe534 of the N-domain, while the phosphate group interacts with Asp409 and Thr411 (the DKTG motif), Asn789, and Asp790 at the phosphorylation site of the P-domain, in cooperation with a Mg^2+^ ion (Fig. 2B). The AlF_4_^−^-ADP-bound state is similar to the E1-AMP-PCP conformation, but the N- and P-domains are tightly bridged by the interaction through ADP and AlF_4_^−^ (Fig. 2A, B), captured in the phosphoryl-transfer intermediate (E1P-ADP) (fig. S6).

**Figure 2:**
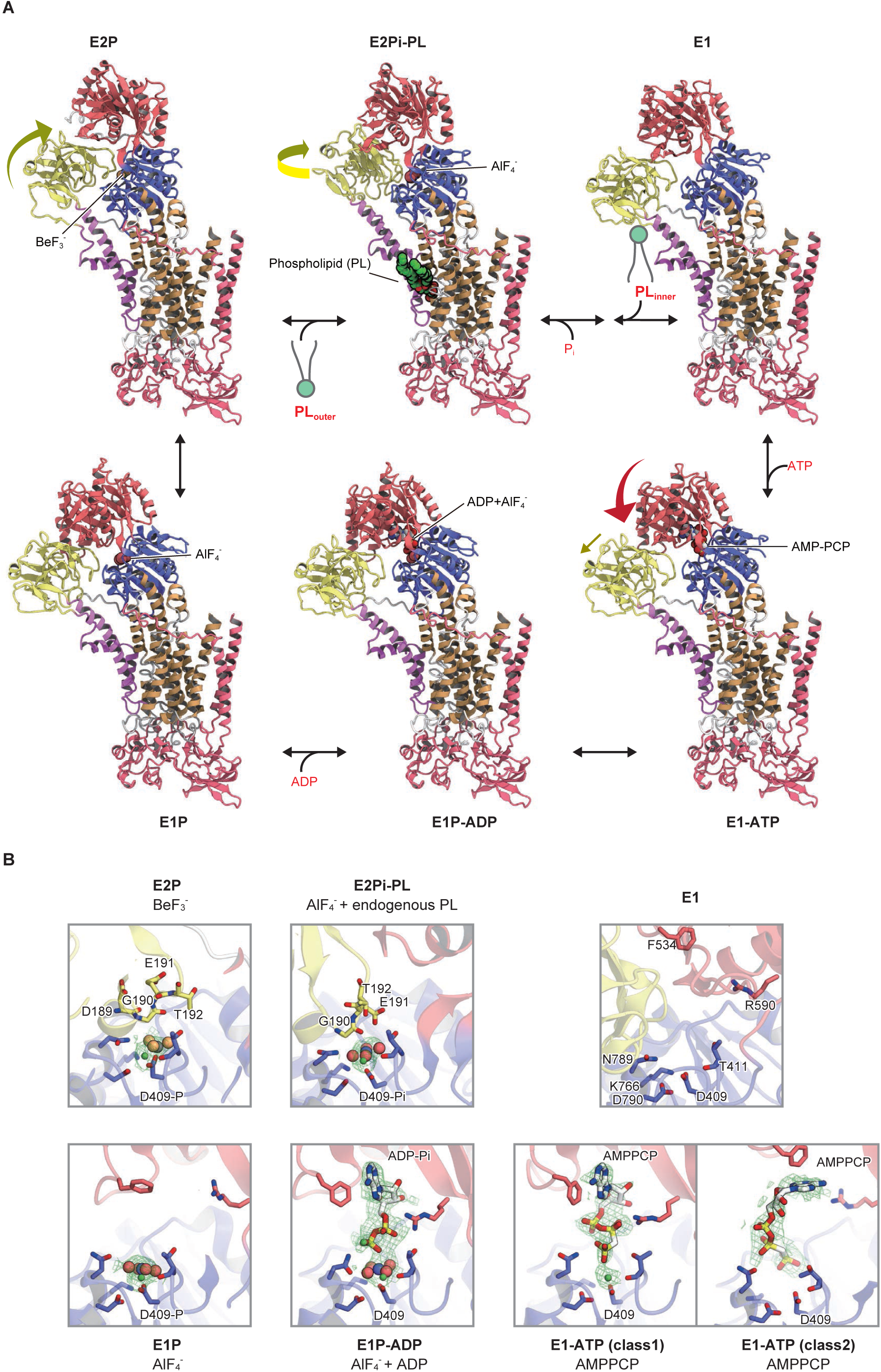
Entire transport cycle of ATP8A1-CDC50a. (**A**) The six different intermediates of ATP8A1-CDC50a during the phospholipid translocation cycle are shown, arranged clockwise as in the Post-Albers reaction cycle: E1, E1-ATP, E1P-ADP, E1P, E2P, and E2Pi-PL. The bound inhibitors are shown in CPK representations. (**B**) Comparison of the phosphorylation sites in each intermediate. AMP-PCP and ADP are shown as sticks, while AlF_4_^−^and BeF_3_^−^ are spheres. Densities are shown as green meshes, contoured at 3.5σ.

Overall, ATP binding and the subsequent phosphoryl transfer reaction induce the proximal arrangement of the N- and P-domains, which is accompanied by a slight outward shift of the A-domain by about 6.5 Å (Fig. 2A, figs. S10-11, and Movie S1). Most notably, the phosphorylation reaction is essentially mediated by the isolated motions of the ATPase domain and does not require any changes in the transmembrane region. The transmembrane segments of ATP8A1 adopt almost the same conformation throughout the transition, which is consistent with the substrate-independent autophorylation of P4-type ATPases (*26*).

The two phosphate analogues, BeF_3_^−^ and AlF_4_^−^, occupy the phosphorylation site in a similar manner, but their coordination geometries are slightly different (Fig. 2A, B). BeF_3_^−^is covalently attached to the carboxylate side chain of Asp409, in coordination with a Mg^2+^ion, and mimics its phosphorylation state. The A-domain is tightly fixed to the phosphorylation site (fig. S7), through the backbone carbonyls of Asp189 and Gly190 in the conserved DGET motif (Fig. 2B). The N-domain is pushed apart from the P-domain and no longer has access to the phosphorylation site, thus representing the ADP-insensitive E2P state (*26*). The particles of the AlF_4_^−^-bound state could be separated into two different classes (fig. S8), and both showed the clear AlF_4_^−^ density at the phosphorylation site. In the first class, the bound AlF_4_^−^ does not mediate any inter-domain interactions, and the catalytic domains adopt a conformation similar to the AlF_4_^−^-ADP bound state (Fig. 2B), likely representing the E1P-like state immediately after the ADP release. In the second class, AlF_4_^−^ mediates the interaction between the N- and A-domains through the DGET motif in a similar manner to the BeF_3_^−^-bound state, but the A-domain is rotated by about 22° around the phosphorylation site, as compared to the BeF_3_^−^-bound state (Fig. 3A). This allows the repositioning of the carboxyl side chain of Glu191 in the DGET motif to provide a catalytic base for the dephosphorylation reaction (Fig. 2B) (*26*), thereby mimicking the dephosphorylation transition-like intermediate (*27*). The rearrangement of the A-domain accompanies the swing-out motion of the TM1-2 segment, which is directly connected to the A domain, consequently creating a large cleft between the M1-2 and M4-5 segments, in which the clear density of a glycero-phospholipid is observed (Fig. 3B). Therefore, the current structure represents the substrate-bound E2Pi state. The rearrangement of the A-domain is likely to be coupled to the binding of the substrate lipid, as it occupies the cleft and pushes out the M1-2 segment (figs. S10 and S11), which explains the substrate-dependent dephosphorylation of P4-ATPases (Fig. 1C) (*26*).

**Figure 3:**
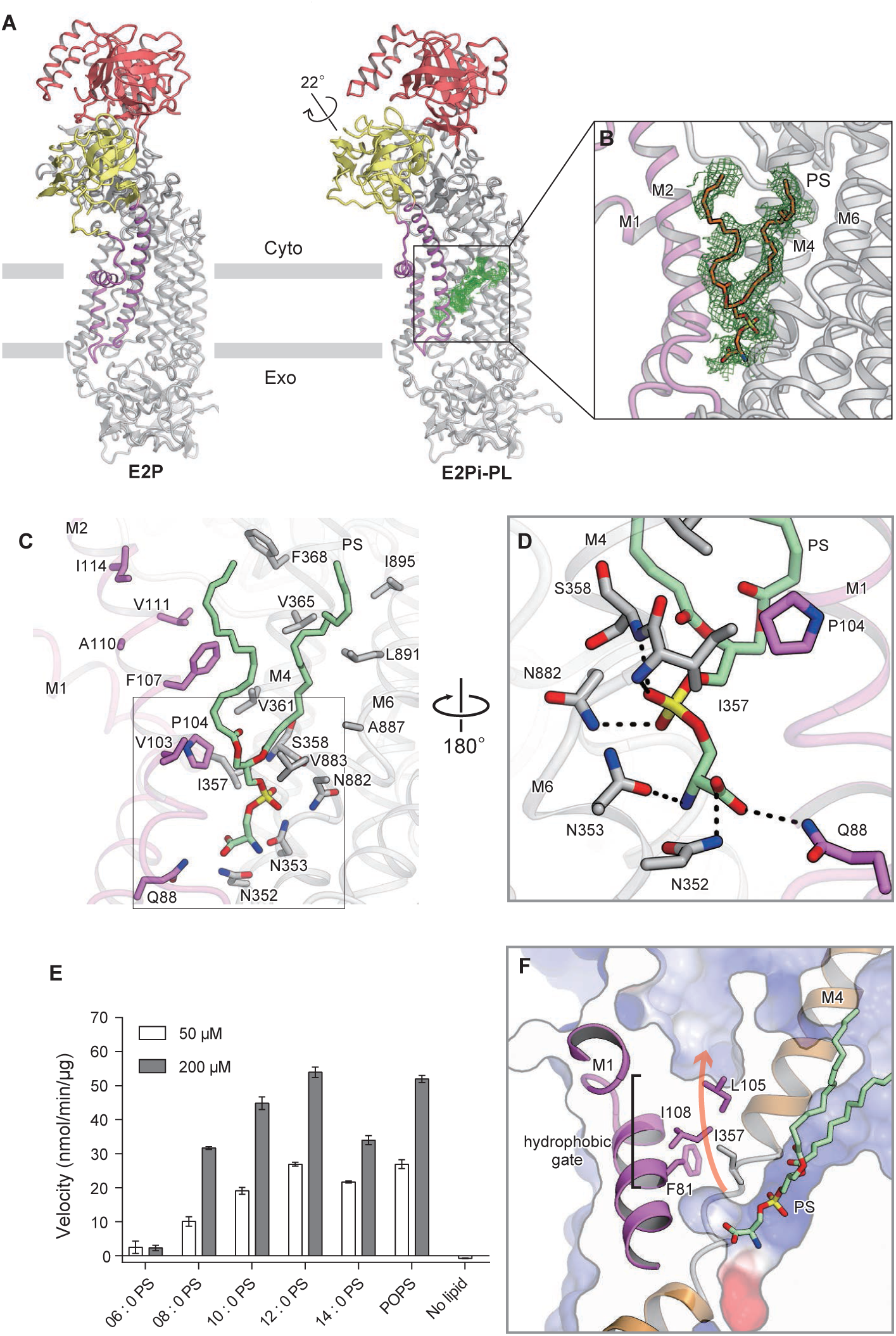
Phospholipid recognition. (**A**) Structural comparison of the E2P and E2Pi-PL states, showing the large rearrangement of M1-2 and the N- and A-domains upon phospholipid binding. (**B**) Cryo-EM density showing the bound endogenous phospholipid (green mesh, 2.5σ). (**C, D**) Phospholipid binding site, viewed parallel to the membrane plane (C), and a close-up view of the head group (D). Residues within 3 Å of the bound phospholipid are shown as sticks. Hydrogen bond interactions are shown as black dashed lines. (**E**) Dependency of the ATPase activity of ATP8A1-CDC50a on the acyl chain length of the PS lipids. Values are mean ± SEM. n = 3. (**F**) Residues constituting the hydrophobic gate are shown. The putative translocation pathway is indicated by an orange arrow.

### Phospholipid recognition

Since the substrate lipids (such as PS or PE) were not added during the purification, it is likely that the endogenous phospholipid contained in the GDN micelles is specifically bound to ATP8A1 in the AlF_4_^−^-bound state (Fig. 3, B and C). ATP8A1 shows PG-, PE-, and PS-dependent ATPase activities, with the highest preference for PS (*12, 16*), and the size and shape of the head group density are in good agreement with those of the serine moiety (fig. S1E). Therefore, we modeled PS into the density. PS is recognized within the open cleft, in which the phosphate group is coordinated by the backbone amide groups of Ile357 and Ser358 in the conserved PISL motif at the unwound kink of M4 and further stabilized by the Asn353 and Asn882 side chains (Fig. 3D), while the attached acyl chains are exposed to the bulk lipid environment and partly accommodated in the hydrophobic pocket formed by the conserved residues in TM2 and TM4, such as Val103, Pro104, Phe107 (M2), Val361, Val365 (M4), Val883, and L891 (M6) (Fig. 3C). In the current cryo-EM map, the acyl chains are most visible near the attached glycerol moiety, and PS molecules with shorter acyl chains showed weaker ATPase activity (Fig. 3E), indicating that the acyl chains, as well as the hydrophilic head group, are specifically recognized in the substrate binding pocket of ATP8A1.

The head group of PS is situated within a small cavity on the extracellular half of the cleft, and is surrounded by hydrophilic residues, such as Gln88, Asn352, Asn353, and Asn882 (Fig. 3D), with which the serine moiety forms hydrogen bonding interactions. Consistently, mutational studies have shown the importance of the uncharged polar residues, Gln88, Gln89, Asn352, and Asn353, for the PS selectivity (*21*–*23*). Such interactions explain the head group preferences of ATP8A1, which has weak selectivities for PE and PG, with head groups that can form similar hydrogen bonding interactions, and no selectivity for PC, with head methyl groups that cannot form such hydrogen bonding interactions. The PC selective P4-ATPases have non-polar residues, such as Ala and Gly, at the corresponding positions (fig. S2), also supporting the notion that the residues constituting this exoplasmic cavity primarily define the head group selectivity.

### Lipid translocation pathway of ATP8A1

In the P2-ATPases, the cytoplasmic ATPase domains are highly associated with the rearrangement of the core TM helices that constitute the cation binding sites (Fig. 4A). Especially, Glu309 of the conserved PEGL motif, located in the unwound M4 kink, constitutes part of the ion binding sites, enabling the coupling between the ion binding and release and the rearrangement in the ATPase domain. In ATP8A1, the M4 segment is similarly kinked at the PISL motif, but the ion binding sites are collapsed by the substitution with hydrophobic residues (Fig. 4B) (*28, 29*). Although the PS binding site partially overlaps with the Ca^2+^ binding site in SERCA (site-II), the arrangement of the surrounding residues remains almost unchanged throughout the transport cycle. The P4-ATPases have been suggested to utilize a different translocation pathway for a large lipid substrate (*8*), and according to the previous mutation studies on the yeast and bovine P4-ATPases, the residues associated with the head group selectivity are mapped along the hydrophilic cleft between the M1-2 and M3-4 segments, highlighting the head group translocation pathway of the P4-ATPase (fig. S12A). In the current PS-bound structure, the head group enters from the exoplasmic leaflet and stays occluded in the middle of the pathway by the side chain of Ile357 in the PISL motif. The mutation of Ile357 to bulky residues, such as Met and Phe, drastically reduced the lipid transport activity, while the mutations to smaller residues only moderately affected the transport activity (*21*), suggesting that Ile357 constitutes a central hydrophobic gate for the lipid translocation, together with other residues on the M1-2 segment, such as Phe81 and Ile105 (Fig. 3F). Since the exoplasmic end is also closed by the M1-2 loop (fig. S12A), the current structure probably represents a partially occluded state. Although PS binding induces a slight reorientation of the Ile357 side chain toward the M1-2 segments (Fig. 4B), the translocation of the hydrophilic head group requires the further rearrangement of the central gate residues, which is probably coupled with the phosphate release from the P-domain. By analogy to the E2P to E2 transition in SERCA, the phosphate release “unlocks” the A-domain and allows a further outward shift of the M1-2 segment, thus inducing the opening of the central hydrophobic gate (*30*). It should be noted that the previous structures of P2-ATPases revealed several lipid binding sites; especially, the E2 structure of SERCA stabilized by thapsigargin and an inhibitor, BHQ (*31*), showed PE binding between the M2 and M4 segments at the intracellular leaflet, corresponding to the putative exit of the lipid translocation pathway in ATP8A1. Furthermore, phospholipids are associated with the positively charged residues at the protein-lipid interface and interplay with the protein during the transport cycle in SERCA (*32*). The ATP8A1-CDC50A complex has clusters of positively charged residues at both the entrance and exit of the translocation pathway (fig. S12B). Therefore, these residues may play important roles in lipid translocation.

**Figure 4:**
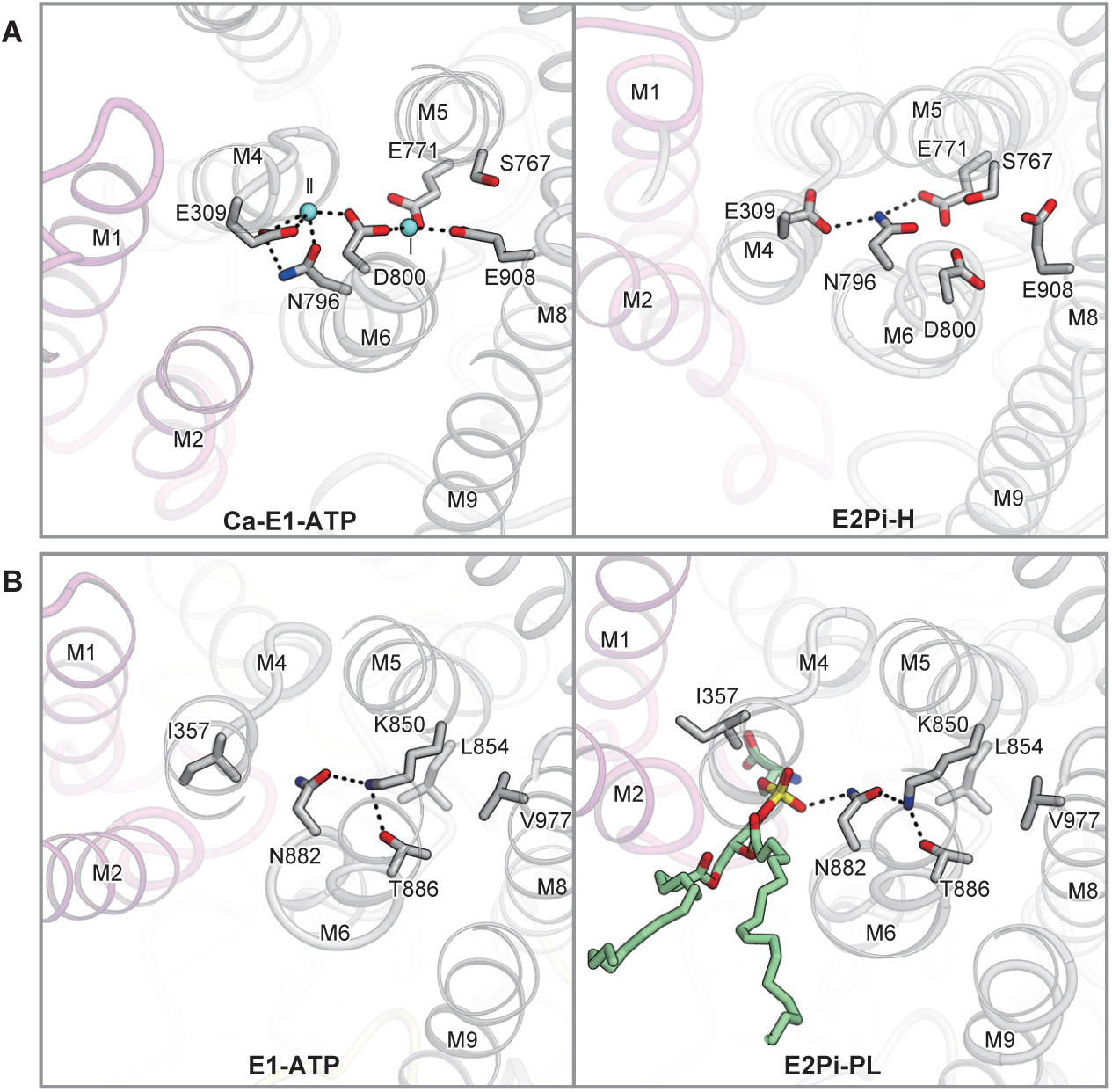
Comparison of the phospholipid binding sites. (**A**) Ca^2+^ binding site of SERCA in the H^+^ binding state (left: PDB 3B9R) and the Ca^2+^ binding state (right: PDB 1T5S), viewed from the cytoplasmic side. Residues involved in Ca^2+^ and H^+^ transport are shown as ball-stick representations. Hydrogen bonds are shown as black dashed lines and the bound Ca^2+^ are pale blue spheres. (**B**) Phospholipid binding site of ATP8A1 in the phospholipid-bound state (left: E2Pi-PL) and the unbound state (right: E1-ATP) from the same viewpoint as in (A). Residues involved in phospholipid translocation and other residues corresponding to those coordinating H^+^ and Ca^2+^ in SERCA are shown as ball-stick representations. Hydrogen bonds are shown as black dashed lines.

### C-terminal auto-regulatory domain

In the BeF_3_^−^-stabilized E2P state, we observed an extra density extending through the cytoplasmic catalytic domains (Fig. 5, A and B), which we assigned as the C-terminal auto-regulatory domain (residues 1117–1140) (*33, 34*), consisting of the conserved GYAFS motif (residues 1119–1123) and a short helical domain (residues 1131–1137), although the ~50-amino acid linker connected to the M10 helix was disordered. The regulatory domain essentially interacts with the N-domain, and the GYAFS motif is specifically recognized by a short loop region (residues 533–540) (Fig. 5B). Notably, Phe1122 occupies the ATP binding site and stacks with Phe534 of the N-domain. The densities of the C-terminal residues are only visible in the BeF_3_^−^-stabilized E2P conformation, and are completely disordered in the other conformations, including the ligand-unbound E1 state. Therefore, the regulatory domain specifically stabilizes ATP8A1 in the E2P conformation, in which the N-domain is somewhat farther apart (Fig. 5C and fig. S10).

**Figure 5:**
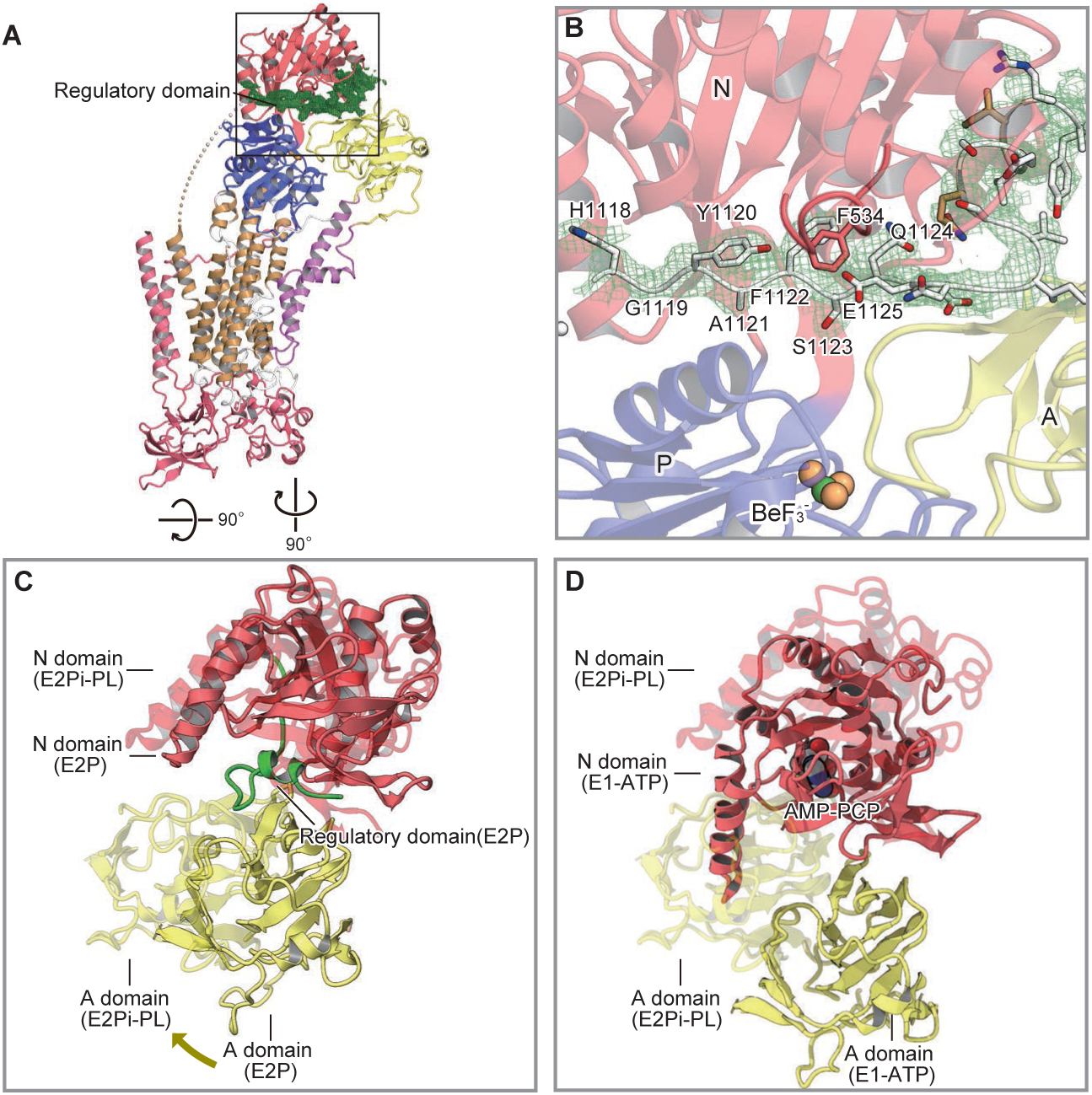
ATP8A1 autoregulation by the C-terminal domain. (**A**) In the BeF_3_^−^-stabilized E2P state, an extra density is observed around the cytoplasmic catalytic domains, corresponding to the C-terminal autoregulatory domain. The density is shown as a green mesh, contoured at 3σ. (**B**) Close up view of the interaction between the N-domain and the regulatory domain. An atomic model of the GYAFS motif and a short helical region in the regulatory domain are modeled into the density. (**C, D**) Arrangements of the N- and A-domains and the regulatory domain, shown for the E2P (C) and E1-ATP (D) states, viewed from the cytoplasmic side. The N- and A-domains of the E2Pi-PL state are superimposed in a transparency representation. The regulatory domain keeps the N-domain apart from the A-domain and thus facilitates the rotational movement of the A-domain around the phosphorylation site in the E2P state (C), while the similar rearrangement is hindered by the N-domain in the E1-ATP state (D).

The C-terminal regulatory domain has different effects between the yeast and mammalian P4-ATPases. In the yeast Drs2p flippase, it essentially exerts an auto-inhibitory effect on the ATPase activity (*35*). However, in the mammalian ATP8A2 flippase, it mediates a rather complicated regulation mode. The partial truncation of the GYAFS motif and the short helical domain results in decreased ATPase activity, whereas the complete loss of the C-terminal residues, including the disordered loop region, restores the ATPase activity to the same level as the wild type enzyme (*36*), indicating that the GYAFS motif and the short helical domain observed in the current cryo-EM map positively modulate the enzymatic reaction. We hypothesize that the regulatory domain keeps the N-domain apart from the A-domain in the E2P state and thus facilitates the rotational rearrangement of the A-domain that is required for PS binding, as the N-domain in the E1/E1-ATP states clashes with this rotational motion (Fig. 5D).

### Unique mechanism of the P4-type ATPase

In this study, we have reported the cryo-EM structures of six different intermediates of ATP8A1; namely, E1, E1-ATP, E1P-ADP, E1P, E2P, and E2Pi-PL, demonstrating the entire transport cycle of the lipid flippase reaction (Fig. 6). While ATP8A1 shares the similar catalytic reaction of ATP hydrolysis with the ion-transporting P2-type ATPases, such as SERCA (*28, 29*), Na^+^/K^+^-ATPase (*37*), and H^+^/K^+^-ATPase (*38*), there are significant differences in their transport mechanisms.

**Figure 6:**
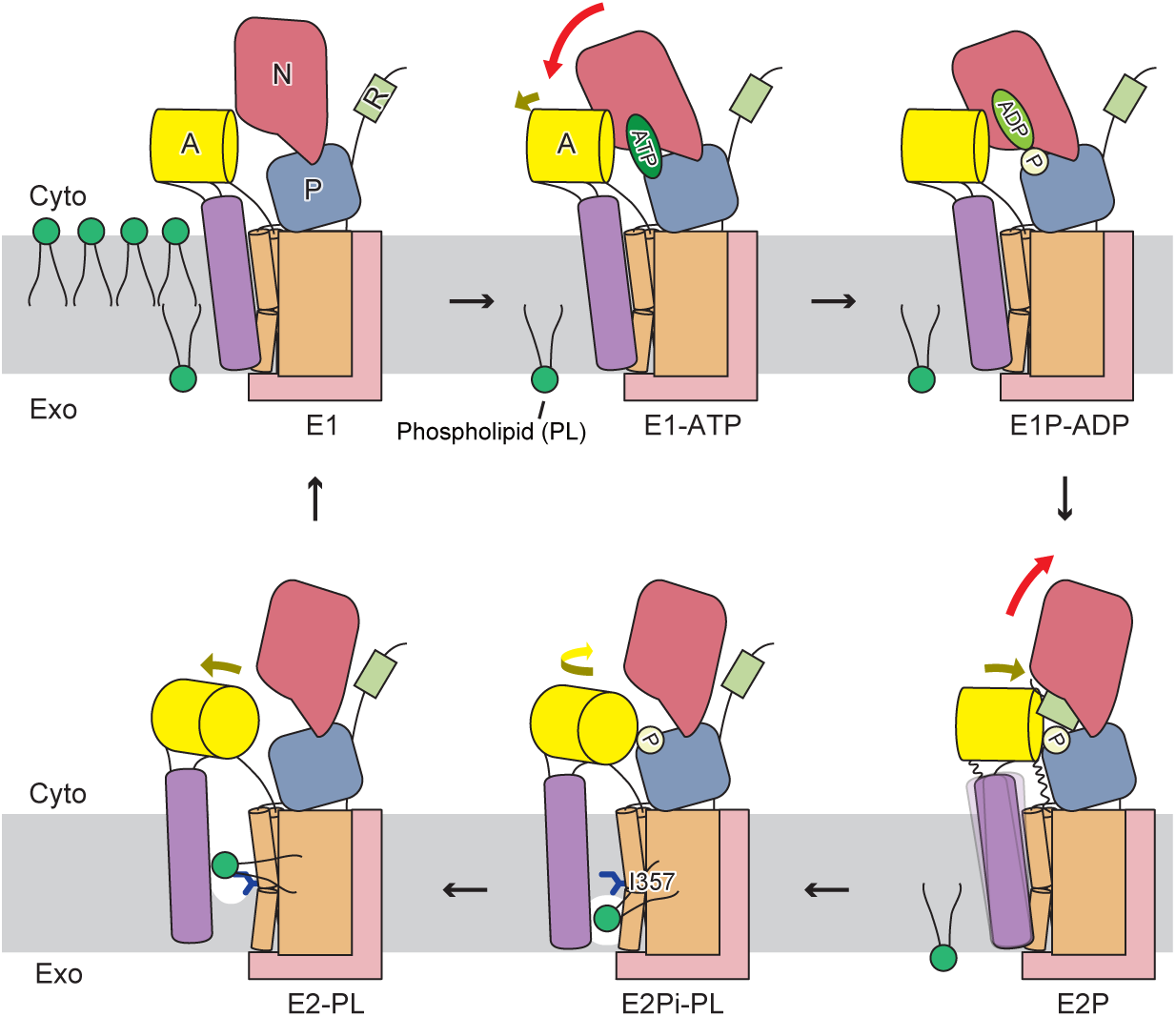
Proposed mechanism of phospholipid translocation. Schematic model of the phospholipid translocation cycle by ATP8A1-CDC50a, according to the Post-Albers mechanism. The model is depicted with the same color codes as in Fig. 1A. ATP binding induces the proximal arrangement of the N- and P-domains, by bridging these domains and slightly forcing out the A-domain. After the phosphoryl transfer reaction, ADP is released from the N-domain, and the A-domain approaches the N-domain and interacts with it, through the TGES motif, to form the E2P state. The C-terminal regulatory domain penetrates between the P- and N-domains and stabilizes the E2P state. The rearrangement of the A-domain induces the flexibility in the M1-2 segments, thus allowing phospholipid binding at the interface between the M1-2 and bulk TM segments. Phospholipid binding induces further rearrangement of the A-domain, thereby facilitating the dephosphorylation reaction (E2Pi-PL). Ile357 constitutes a hydrophobic gate that occludes the middle of the translocation pathway. Phospholipid translocation to the cytoplasmic leaflet is probably coupled to the phosphate release at the P-domain, allowing the further outward shift of the M1-2 segment (E2-PL). The translocated phospholipid laterally diffuses to the cytoplasmic leaflet, and the enzyme adopts the E1 conformation, ready to initiate another reaction cycle.

ATP8A1 maintains the structural rigidity of the transmembrane region throughout the transport cycle, as the core TM segments (M3–10) could be superimposed well in all of the intermediates (RMSD = 0.294 ~ 0.689). The critical rearrangement in SERCA occurs during the E1P to E2P transition, in which the A-domain rearrangement toward the phosphorylation site induces the opening of the “luminal gate” composed of the M1–4 segments and alters the affinity for the substrate ions (fig. S13) (*29*). While the A-domain of ATP8A1 undergoes a similar rearrangement during this transition, the conformational change is limited to the region proximal to the A-domain, and the luminal side remains unchanged (fig. S13). This rigidity is probably achieved by the tight association with CDC50a, which holds the M3–10 segments of ATP8A1 on both the luminal and cytoplasmic sides. Most notably, the loop connecting M3-4 and the cytoplasmic end of M4 are held by the interaction with CDC50a, which contributes to the stability of the M3 and M4 segments (figs. S3D, E and S13B). The deletion of the CDC50a N-terminal tail, which interacts with the cytoplasmic end of the M4 segment, decreased the flippase activities of P4-ATPases (*39, 40*). Therefore, the rigidity of the TM segments is important for the transport activity of P4-ATPase, and consequently, the lipid translocation is essentially accomplished by the mobile segments of M1-2. While the phosphorylation-induced A-domain rearrangement in ATP8A1 itself causes only minor changes on the luminal side, the density of the M1-2 segment near the A-domain is more disordered in the E2P conformation (fig. S14), suggesting the higher flexibility in the linker region. Such flexibility may facilitate the subsequent binding of the phospholipid between the M1-2 and M3-4 segments by allowing the swing-out motion of the M1-2 segment, as observed in the AlF_4_^−^-stabilized dephosphorylation transition-like state. Overall, the P4-ATPases have evolved a distinct translocation mechanism and translocation pathway, while sharing the similar rearrangement of the cytoplasmic domains with the canonical ion-transporting P-type ATPases. Therefore, the current study highlights the diverse mechanisms adopted by the P-type ATPase family proteins.

## Acknowledgments

We thank Hisato Hirano for assistance in generating the movie, Dr. Hiroshi Nishimasu for fruitful discussions, Dr. Takanori Nakane for assistance with the single particle analysis, and the Structural Biophysics team at Mitsubishi Tanabe Pharma Corporation, especially Dr. Toshiyuki Kishida, for technical advice about model building. We also thank the staff scientists at the Cryo-EM facility in The University of Tokyo, especially Dr. Kan Kobayashi, Dr. Tsukasa Kusakizako, Dr. Haruaki Yanagisawa, Dr. Akihisa Tsutsumi, Dr. Masahide Kikkawa, and Dr. Radostin Danev.

## Author contributions

M.H. prepared the cryo-EM samples and performed the functional analyses. M.H. and T.N. collected and processed the cryo-EM data, and built the structures. K.Y. assisted data processing and structure refinement. M.H., T.N., and O.N. wrote the manuscript. T.N. and O.N. supervised the research.

## Data and materials availability

Cryo-EM density maps have been deposited in the Electron Microscopy Data Bank, under the accession codes EMD-9931 (E1 class1), EMD-9932 (E1 class2), EMD-9933 (E1 class3YY), EMD-9935 (E1-ATP class1), EMD-9934 (E1-ATP class2), EMD-9936 (E1-ATP class3), EMD-9937 (E1P-ADP), EMD-9938 (E2P class1), EMD-9939 (E2P class2), EMD-9940 (E2P class3), EMD-9941 (E2Pi-PL), and EMD-9942 (E1P). Atomic coordinates have been deposited in the Protein Data Bank, under the accession codes 6K7G (E1 class1), 6K7H (E1 class2), 6K7J (E1-ATP class1), 6K7I (E1-ATP class2), 6K7K (E1P-ADP), 6K7L (E2P-class2), 6K7M (E2Pi-PL), and 6K7N (6K7N).

