## Supplemental Figure, Table, Methods for "Cryo-EM structures capturing the entire transport cycle of the P4-ATPase flippase"

**This PDF file includes:**

Materials and Methods
Figs. S1 to S14
Table S1
Captions for Movie S1

**Other Supplementary Materials for this manuscript include the following:**

Movie S1

#### Materials and Methods

Expression and purification of the ATP8A1-CDC50a heterodimer

The human ATP8A1 (Uniprot: Q9Y2Q0-2) cDNA was purchased from Kazusa DNA Research Institute. The human cell division control protein 50 A (CDC50A, Uniprot: Q9NV96) cDNA was synthesized and codon-optimized for expression in human cell lines. Both cDNAs were cloned into the pcDNA3.4 vector. The ATP8A1 sequence was fused with an N-terminal His8 tag and enhanced green fluorescent protein (EGFP), followed by a tobacco etch mosaic virus (TEV) protease site.

HEK293F cells were grown and maintained in FreeStyle 293 medium (Gibco) at 37°C, with 8% $CO_2$ under humidified conditions. Cells were not tested for mycoplasma contamination. Cells were transiently transfected at a density of $2.0 \times 10^6$ cells $ml^{-1}$, with the plasmids and FectoPRO (Polyplus). Approximately 270 μg of the ATP8A1 plasmid and 90 μg of the CDC50a plasmid were premixed with 360 μL FecroPRO reagent in 80 mL of fresh FreeStyle 293 medium, for 10–20 min before transfection. For transfection, 80 ml of the mixture was added to 0.8 L of cell culture and incubated at 37°C in the presence of 8% $CO_2$. After 18–20 h, 2.2 mM valproic acid was added, and the cells were further incubated at 30 °C in the presence of 8% $CO_2$ for 48 h before collection. The cells were collected by centrifugation (3,000 × g, 10 min, 4 °C) and disrupted by dounce homogenization in hypotonic buffer (50 mM HEPES-NaOH (pH 7.5), 10 mM KCl, 0.04 mg $ml^{-1}$ DNase I, and protease inhibitor cocktail). The membrane fraction was collected by ultracentrifugation (138,000 × g, 1 h, 4 °C), and solubilized for 1 h at 4 °C in buffer (50 mM HEPES-NaOH (pH 7.5), 300 mM NaCl, 1.5% (w/v) *N*-dodecyl β-d-maltoside (DDM), and 0.15% (w/v) cholesterol hemisuccinate (CHS)). After ultracentrifugation (138, 000 × g, 30 min, 4 °C), the supernatant was incubated with AffiGel 10 (Bio-Rad) coupled with a GFP-binding nanobody (*41*), and incubated for 2 h at 4 °C. The resin was washed five times with 3 CVs of wash buffer (50 mM HEPES-NaOH (pH 7.5), 300 mM NaCl, and 0.06% GDN), and gently suspended overnight with TEV protease to cleavage the His8-EGFP tag. After the TEV protease digestion, the flow-through was pooled, concentrated, and purified by size-exclusion chromatography on a Superose 6 Increase 10/300 GL column (GE Healthcare), equilibrated with SEC buffer (20 mM HEPES-NaOH (pH 7.5), 150 mM NaCl, and 0.06% GDN). The peak fractions were pooled, concentrated to 4–8 mg $ml^{-1}$, and kept at -80°C before use.

Lipid-dependent ATPase activity assay

All ATPase activity assays were performed using BIOMOL® Green Reagent (Enzo Life Sciences Inc.), with modifications from the previously described protocols (*12*, *16*). A 2 ng portion of the purified ATP8A1-CDC50a was incubated in a 50 μl reaction volume, containing 50 mM HEPES-NaOH (pH 7.5), 150 mM NaCl, 1 mM ATP, 0.03% GDN, 0.2% sodium cholate, 12.5 mM

$MgCl_2$, 1 mM DTT, and the respective lipids at the indicated concentrations. Reactions were performed at 37°C for 25 min, and terminated by the addition of the reagent from the kit. After the mixture was incubated for 25 min at room temperature, the amount of released inorganic phosphate was determined colorimetrically, using a microplate reader (Synergy H1) or a Nanodrop spectrophotometer, by measuring the absorbance at 620 nm. The amount of released phosphate was determined by calibration with known concentrations of the phosphate standard (BML-KI102). All data points represent at least triplicate analyses, and each experiment was repeated at least twice independently. Data were analyzed with GraphPad Prism 6.

Electron microscopy sample preparation

The purified protein solution of ATP8A1-CDC50a was mixed with inhibitor solutions, with the following final concentrations: E1-ATP, 5 mM $MgCl_2$ and 2 mM AMP-PCP; E1P-ADP, 5 mM $MgCl_2$, 5 mM NaF, 1 mM $AlCl_3$, and 5 mM ADP; E2P, 5 mM $MgCl_2$, 10 mM NaF, and 2 mM $BeSO_4$; E1P and E2Pi-PL, 5 mM $MgCl_2$, 10 mM NaF, and 2 mM $AlCl_3$. After an incubation for 1 hour on ice, the protein solutions were applied to a freshly glow-discharged Quantifoil holey carbon grid (R1.2/1.3, Cu/Rh, 300 mesh), using a Vitrobot Mark IV (FEI) at 4°C with a blotting time of 4s under 99% humidity conditions, and then the grids were plunge-frozen in liquid ethane.

Electron microscopy data collection and processing

The prepared grids were transferred to a Titan Krios G3i microscope (Thermo Fischer Scientific), running at 300 kV and equipped with a Gatan Quantum-LS Energy Filter (GIF) and a Gatan K3 Summit direct electron detector in the electron counting mode. Imaging was performed at a nominal magnification of 105,000×, corresponding to a calibrated pixel size of 0.83 Å/pix (The University of Tokyo, Japan). Each movie was recorded for 3.2 seconds and subdivided into 54 frames. The electron flux rate was set to 14 $e^-$/pix/s at the detector, resulting in an accumulated exposure of 64 $e^-/Å^2$ at the specimen. The data were automatically acquired using the SerialEM software (*42*), with a defocus range of -0.8 to -1.6 μm. More than 3,500 movies were acquired for each condition grid, and the numbers of total images are described in Supplementary data table 1. For all datasets, the dose-fractionated movies were subjected to beam-induced motion correction using MotionCor2 (*43*), and the contrast transfer function (CTF) parameters were estimated using CTFFIND4 (*44*).

For the E1 state dataset without inhibitor, 1,417,719 particles were initially picked from the 3,590 micrographs by using the Laplacian-of-Gaussian picking function in RELION-3 (*24*), and extracted with down-sampling to a pixel size of 3.24 Å/pix. These particles were subjected to several rounds of 2D and 3D classifications. The best class contained 702,610 particles, which were then re-extracted with a pixel size of 0.83 Å/pix and subjected to 3D refinement. The resulting 3D model and particle set were subjected to per-particle CTF refinement, beam-tilt refinement, Bayesian polishing (*45*), and 3D refinement. The no-align 3D classification using a mask covering the N- and A-domains resulted in 3 classes of maps. The final 3D refinement and postprocessing of the three classes yielded maps with global resolutions of 3.32 Å (class 1), 3.22 Å (class 2), and 3.42 Å (class 3), according to the FSC = 0.143 criterion (*47*). The local resolution was estimated. As the density maps of the N- and A-domains were confirmed for class 1 and class 2, these maps were used for modeling. The processing strategy is described in Supplementary figure 4.

For the E1-ATP state dataset in the presence of AMP-PCP, 1,405,466 particles were initially picked from the 4,333 micrographs, and extracted with a pixel size of 3.24 Å/pix, as described above. These particles were subjected to two rounds of 3D classifications. The best class from the 3D classification contained 756,868 particles, which were then re-extracted with a pixel size of 0.83 Å/pix and subjected to 3D refinement. The resulting 3D model and particle set were subjected to per-particle CTF refinement, beam-tilt refinement, Bayesian polishing, and 3D refinement, and 3D classification. A subset of 753,281 particles was used to focus on the A- and N-domains by the no-align 3D classification using a mask covering the N- and A-domains resulted in 4 classes of maps. The final 3D refinement and postprocessing of three classes, in which clear densities were confirmed among the four classes, yielded maps with global resolutions of 3.08 Å (class 1), 3.22 Å (class 2), and 3.32 Å (class 3), according to the FSC = 0.143 criterion. The local resolution was estimated. As the density maps of the N- and A-domains were confirmed for class 1 and class 2, these maps were used for modeling. The processing strategy is described in Supplementary figure 5.

For the E1P-ADP state dataset in the presence of $AlF_4^-$ and ADP, 1,097,443 particles were initially picked from the 4,395 micrographs, and extracted with down-sampling to a pixel size of 3.24 Å/pix, as described above. These particles were subjected to three rounds of 3D classifications. The best class from the 3D classification contained 316,665 particles, which were then re-extracted with a pixel size of 0.83 Å/pix and subjected to 3D refinement, using a soft mask covering the proteins and micelle. The resulting 3D model and particle set were subjected to per-particle CTF refinement, beam-tilt refinement, and Bayesian polishing before the final 3D refinement and postprocessing, yielding a map with a global resolution of 3.04 Å according to the Fourier shell correlation (FSC) = 0.143 criterion. Finally, the local resolution was estimated using RELION-3. The processing strategy is described in Supplementary figure 6.

For the E2P state dataset in the presence of $BeF_3^-$, 2,315,815 particles were initially picked from the 6,257 micrographs, and extracted with a pixel size of 3.32 Å/pix as described above. These particles were subjected to several rounds of 3D classifications. The best class from the 3D classification contained 855,936 particles, which were then re-extracted with a pixel size of 1.83 Å/pix and subjected to 3D refinement. The resulting 3D model and particle set were resized to 0.83

Å/pix and subjected to per-particle CTF refinement, beam-tilt refinement, Bayesian polishing, and 3D refinement. The 3D classification of the resulting particles resulted in 4 classes of maps. The final 3D refinement and postprocessing of three classes, in which clear signals were found among the four classes, yielded maps with global resolutions of 2.80 Å (class 1), 2.83 Å (class 2), and 2.66 Å (class 3), according to the FSC = 0.143 criterion. The local resolution was estimated. As the density maps of the N- and A-domains were confirmed for class 2, this map was used for modeling. The processing strategy is described in Supplementary figure 7.

For the E2Pi-PL and E1P state datasets in the presence of $AlF_4^-$, 2,203,101 particles were initially picked from the 6,843 micrographs, and extracted with a pixel size of 3.24 Å/pix as described above. These particles were subjected to several rounds of 3D classification and divided into three classes. The two classes in which clear densities were confirmed were separately re-extracted with a pixel size of 0.83 Å/pix and subjected to 3D refinement. The resulting 3D model and particle set were subjected to per-particle CTF refinement, beam-tilt refinement, Bayesian polishing 3D refinement. Postprocessing of two classes yielded maps with global resolutions of 2.95 Å (E2Pi-PL) and 2.84 Å (E1P), according to the FSC = 0.143 criterion. The local resolution was estimated, and the processing strategy is described in Supplementary figure 8.

<u>Model building and validation</u>

The density maps of the E1P-ADP and E2P states of the ATP8A1-CDC50a complex exhibited sufficient quality for *de novo* model building manually in COOT (fig. S9) (*48*), facilitated by the previous crystal structures of SERCA (PDB 3B9B, 3B9R). After manual adjustment of the models, structure refinement was performed with 'phenix.real_space_refine' (*49*) against the working map in PHENIX (*50*). The atomic models of the other states were built by using the E1P-ADP model as the starting model. Manual real-space refinement in COOT was initially used to adjust all transmembrane helices and side chains to their approximate locations in the experimental map. For the E1 state, the cytoplasmic N- and A-domains were flexible, and the N- and A-domains were modeled into the map by rigid-body fitting and manually modified in COOT. The residue sequences (1–34, 434–442, 698–723, 1064–1116 and 1141–1149) of ATP8A1 and (1–26 and 352–361) of CDC50a were omitted, as the corresponding densities were not visible in all of the maps. The densities corresponding to residues (45–56 and 273–280) were not visible in the E2P state.

The potential overfitting of the refined models was tested by using a cross-validation method, as described previously (*51*). Briefly, the final models were 'shaken' by introducing random shifts to the atomic coordinates up to 0.5 Å, and were refined against the first half map. These shaken-refined models were used to calculate the FSC against the same first half maps (FSChalf1 or work), and the second half maps (FSChalf2 or free) that were not used for the refinement, using phenix.mtriage (*52*). The small differences between the FSChalf1 and FSChalf2 curves indicated no severe overfitting of the models. The curves representing model vs. full map were calculated, based on the final model and the full, filtered and sharpened map. The statistics of the 3D reconstruction and model refinement are summarized in Supplementary Data Table 1. All molecular graphics figures were prepared with CueMol (http://www.cuemol.org) and UCSF Chimera (*53*).

**Fig. S1. Biochemical characterization of the ATP8A1-CDC50a complex.**

(**A**) Representative size-exclusion chromatography profile of ATP8A1-CDC50a. (**B**) SDS-PAGE analysis of the ATP8A1-CDC50a peak fractions in the SEC purification. (**C**) GFP-FSEC profiles for GFP-tagged ATP8A1, expressed with CDC50a (red) and ATP8A1 alone (blue). The arrows indicate the elution positions of the void volume, the ATP8A1-CDC50a complex, and the free EGFP. (**D**) Aluminum fluoride ($AlF_4^-$) and beryllium fluoride ($BeF_3^-$) inhibition of the ATPase activity of ATP8A1-CDC50a in GDN micelles, stimulated with 200 μM POPS. Each point represents the mean ± SEM of three separate measurements. (**E**) Chemical structures of phospholipids. Abbreviations: PS phosphatidylserine, PE phosphatidylethanolamine, PA phosphatidic acid, PC phosphatidylcholine, and PG phosphatidylglycerol. The hydrophilic head groups attached to the basic phospholipid structures are shown in red.

**Fig. S2. Sequence alignment of the human and yeast P4 ATPases and the major P2 ATPases.**

Sequence alignment of five PS selective P4-ATPases: human ATP8A1 (UniProt: Q9Y2Q0-2), human ATP8A2 (UniProt: Q9NTI2), bovine ATP8A2 (UniProt: C7EXK4), human ATP9A (UniProt: O75110), and *Saccharomyces cerevisiae* Drs2 (UniProt: P39524); three PC selective P4 ATPases: human ATP8B1 (UniProt: O43520), human ATP10A (UniProt: O60312), and *Saccharomyces cerevisiae* Dnf1 (UniProt: P32660), and two major P2 ATPases: shark $Na^+/K^+$-ATPase (UniProt: Q4H132) and sarco (endo) plasmic reticulum $Ca^{2+}$-ATPase (SERCA) isoform 1a (UniProt: P04191). The conserved domains and transmembrane helices of hATP8A1 are indicated above the sequences. The conserved QQ motif, DGET motif, TGD motif, phosphorylation site, and GYAF motif are indicated by black boxes. The similarly conserved residues are indicated in red letters. The positions of the disease-related mutations of hATP8A2 and hATP8B1 are indicated with cyan boxes. The residues involved in the phospholipid recognition and translocation in the P4-ATPase are highlighted with pink circles above the alignment. The residues involved in the $Ca^{2+}$ and $H^+$ transport in SERCA are highlighted with black circles below the alignment. In ScDrs2, hATP8B1, hATP10A, and ScDnf1, the internal loops are omitted for clarity (shown as wavy lines).

**Fig. S3. Heterodimer interaction of ATP8A1 and CDC50a.**

(**A**) Electrostatic surface potentials of ATP8A1 and CDC50a, displayed as a color gradient from red (negative) to blue (positive). (**B**) Close-up view of the cholesterol binding site between ATP8A1 and CDC50a. Densities are contoured at 3.5σ. (**C**) Glycosylation and disulfide sites of CDC50a. The NAG densities were identified at Asn107 and Asn294. (**D**) Tyr329 and Trp328, at the tip of the M3-4 loop of ATP8A1, interact with hydrophobic residues and the N-glycan attached to Asn180 of CDC50a. (**E**) Interactions between the loop segments of ATP8A1 (yellow) and the N-terminal residues of CDC50a. (**F**) Interactions between M9-10 of ATP8A1 and TM1-2 of CDC50a.

**Fig. S4. Data processing of the E1 state without inhibitor.**

(**A**) Representative cryo-EM image of the ATP8A1-CDC50a complex without inhibitor, recorded on a 300 kV Titan Krios with a K3 camera. (**B**) Data processing workflow of the single particle image processing and the local resolution analysis. Particles were separated into three groups by no-align 3D classification, with the mask covering the N- and P-domains. The applied mask is shown in transparent white. (**C**) Cross-validation FSC curves for map-to-model fitting.

**Fig. S5. Data processing of the E1-ATP state bound to AMP-PCP.**

(**A**) Representative cryo-EM image of the ATP8A1-CDC50a complex in the presence of AMP-PCP. (**B**) Data processing workflow of single particle image processing and local resolution analysis. Particles were separated into four groups by no-align 3D classification, with the mask covering the N- and P-domains, and the three classes were selected for the Refine 3D process. The applied mask is shown in transparent white. (**C**) Cross-validation FSC curves for map-to-model fitting. (**D**) Comparison of the two AMP-PCP bound classes (class 1 and class 2).

**Fig. S6. Data processing of the E1P-ADP state bound to $AlF_4^-$ and ADP.**

(**A**) Representative cryo-EM image of the ATP8A1-CDC50a complex in the presence of $AlF_4^-$ and ADP. (**B**) Data processing workflow of the single particle image processing and the local resolution analysis. (**C**) Cross-validation FSC curves for map-to-model fitting.

**Fig. S7. Data processing of the E2P state bound to $BeF_3^-$.**

(**A**) Representative cryo-EM image of the ATP8A1-CDC50a complex in the presence of $BeF_3^-$. (**B**) Data processing workflow of the single particle image processing and the local resolution analysis. (**C**) Cross-validation FSC curves for map-to-model fitting.

**Fig. S8. Data processing of the E1P and E2Pi-PL states bound to $AlF_4^-$.**

(**A**) Representative cryo-EM image of the ATP8A1-CDC50a complex in the presence of $AlF^{4-}$. (**B**) Data processing workflow of the single particle image processing and the local resolution analysis. The particles were 3D classified into three groups, and two groups were selected for further processing, showing distinct conformations. (**C**) Cross-validation FSC curves for map-to-model fitting.

**Fig. S9. Atomic model of ATP8A1-CDC50a in the density maps.**

(**A**, **B**) Cryo-EM density and atomic model of each segment of ATP8A1 (A) and CDC50a (B) in the E1P-ADP state. All maps are contoured at 3.5σ.

**Fig. S10. Comparison of the transmembrane domains.**

The transmembrane domains of ATP8A1 are shown for the six intermediates, viewed from the cytoplasmic side and arranged clockwise as in the Post-Albers reaction cycle: E1, E1-ATP, E1P-ADP, E1P, E2P, and E2Pi-PL. M1-2 and M3-10 are colored purple and brown, respectively. The transmembrane segments adopt similar conformations during the translocation cycle.

**Fig. S11. Overall comparison of the ATPase domains.**

The ATPase domain of ATP8A1 is shown for the six intermediates, viewed from the cytoplasmic side and arranged clockwise as in the Post-Albers reaction cycle: E1, E1-ATP, E1P-ADP, E1P, E2P, and E2Pi-PL. The A, N, and C-terminal regulatory domains are colored yellow, red, and green, respectively. These domains undergo dynamic conformational changes coupled with the ATP hydrolysis and auto-phosphorylation.

**Fig. S12. Hydrophilic cleft and residues affecting the phospholipid specificity.**

(**A**) Cross-sections of the electrostatic surface potentials in the E2Pi-PL state. The potentials are displayed as a color gradient from red (negative) to blue (positive). Residues associated with the head group selectivity are mapped along the hydrophilic cleft between the M1-2 and M3-4 segments

(*22*, *23*, *54*–*56*), viewed from the membrane side. (**B**) Clusters of positively charged residues at both the entrance and exit of the translocation pathway.

**Fig. S13. Comparison of the movements of M1-4 with substrate binding in SERCA and ATP8A1-CDC50a.**

(**A**) Structural comparison of SERCA between E1P-ADP (cyan, PDB 1T5T) and E2P (orange, PDB 3B9B) (*28*, *29*). SERCA involves the opening of the luminal gate accompanied with the phosphorylation-induced rearrangement of the A-domain. (**B**) Structural comparison of ATP8A1-CDC50a between E1P-ADP and E2P. ATP8A1 undergoes only minor changes on the luminal side during the E1P to E2P transition, and is stabilized by the interaction with CDC50a. The structures are viewed from the membrane side (left) and the cytoplasmic side (right).

**Fig. S14. A-domain rearrangement and the linker disorder upon phosphorylation.**

Rearrangement of the A-domain during the E1P (left) to E2P (right) transition. Density maps for the A-domain and the M1-2 segments are superimposed, with the color according to the local resolution (red: high resolution to blue: low resolution). The dotted lines indicate disordered regions in the E2P state.

**Table S1. Data collection, processing, model refinement, and validation.**

**Movie S1. Entire phospholipid transport cycle of ATP8A1-CDC50a**

Schematic movie of the phospholipid translocation cycle by ATP8A1-CDC50a, according to the Post-Albers mechanism, with the morphed series of the conformational transitions through E1, E1-ATP, E1P-ADP, E1P, E2P, and E2Pi-PL. Each domain is illustrated with the same color code as in Fig. 1.

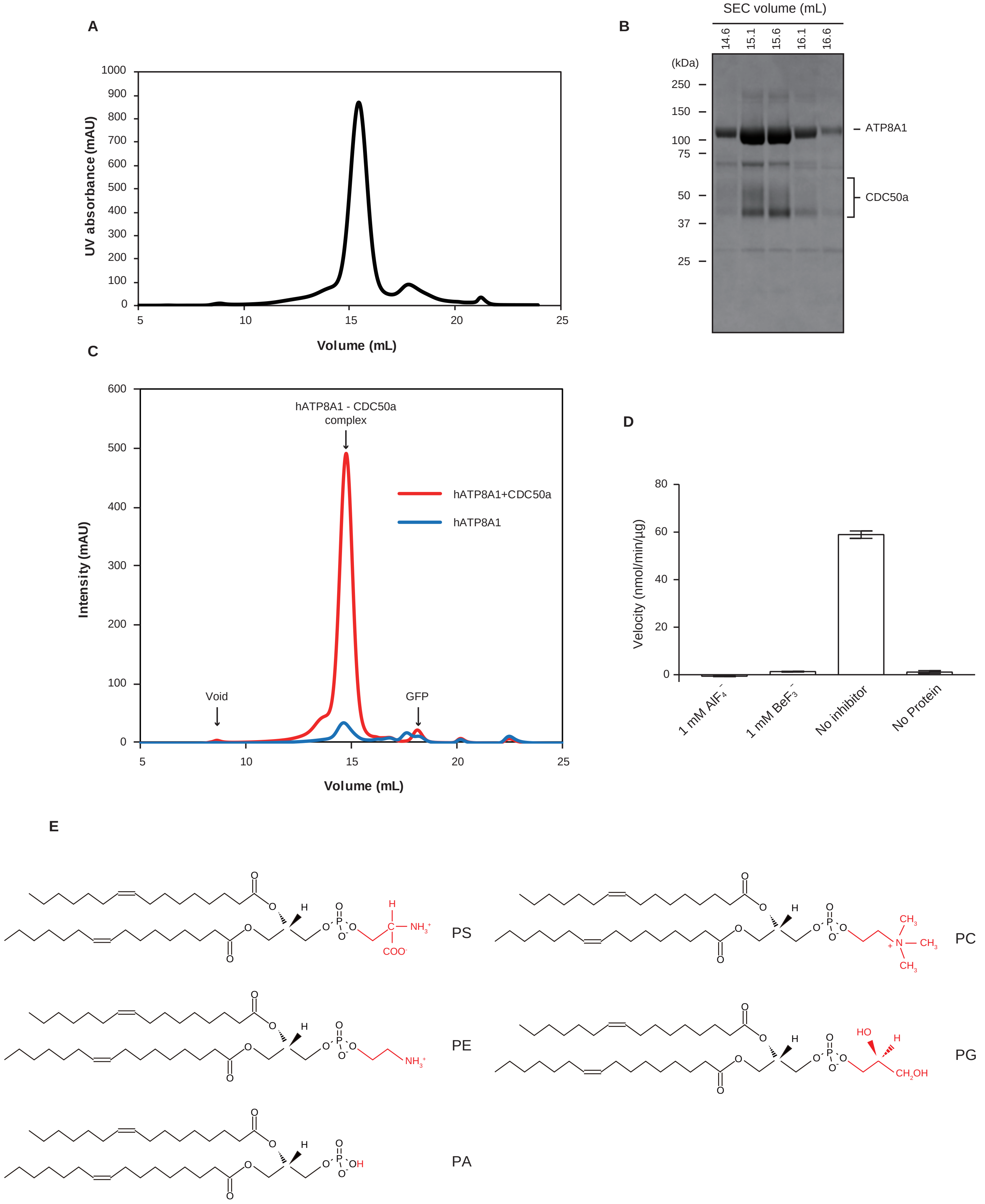


### Supplementary Figure 1

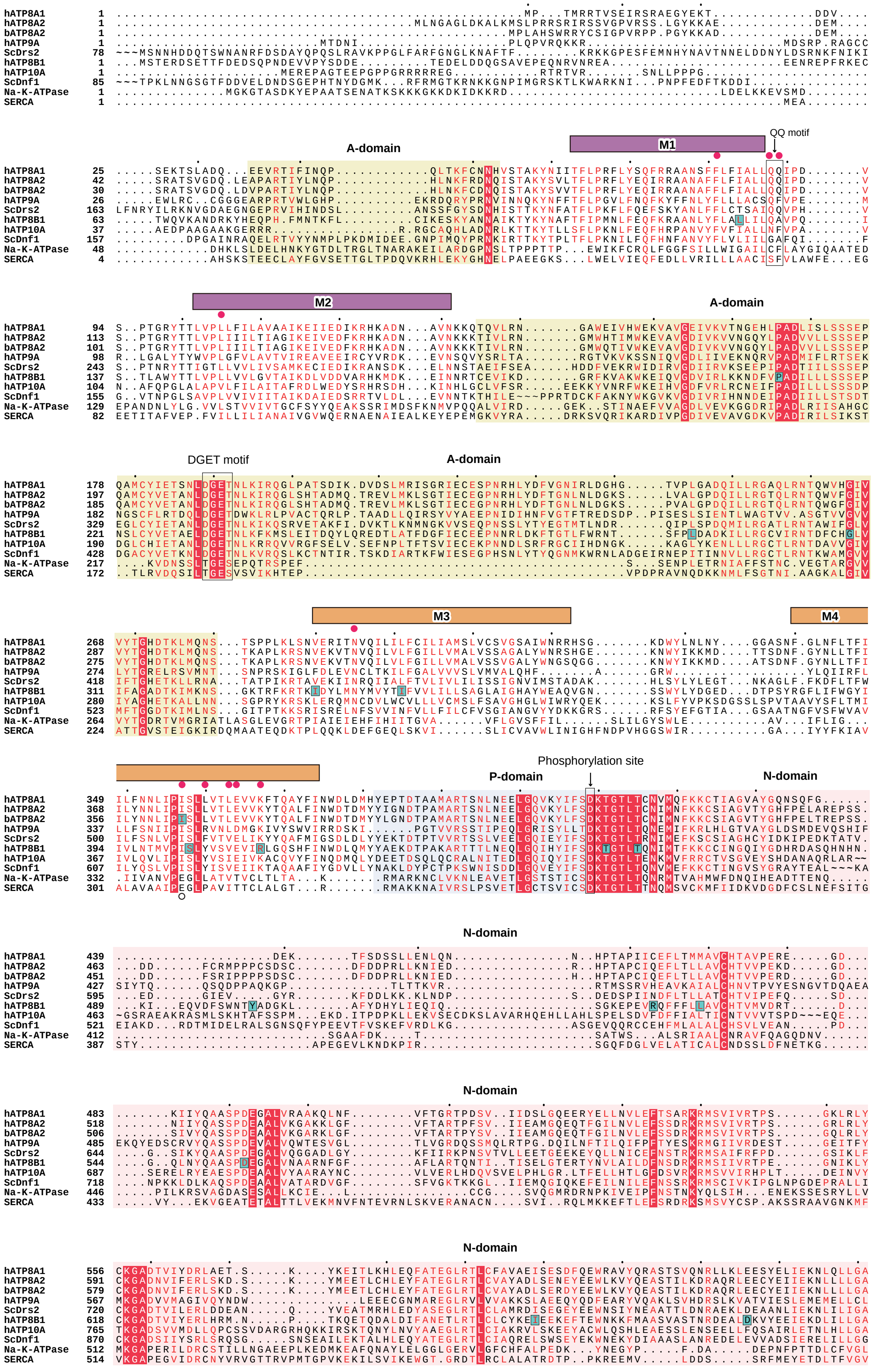
A-domain
M1
QQ motif
M2
DGET motif
M3
M4
Phosphorylation site
P-domain
N-domain
hATP8A1
hATP8A2
bATP8A2
hATP9A
ScDrs2
hATP8B1
hATP10A
ScDnf1
Na-K-ATPase
SERCA

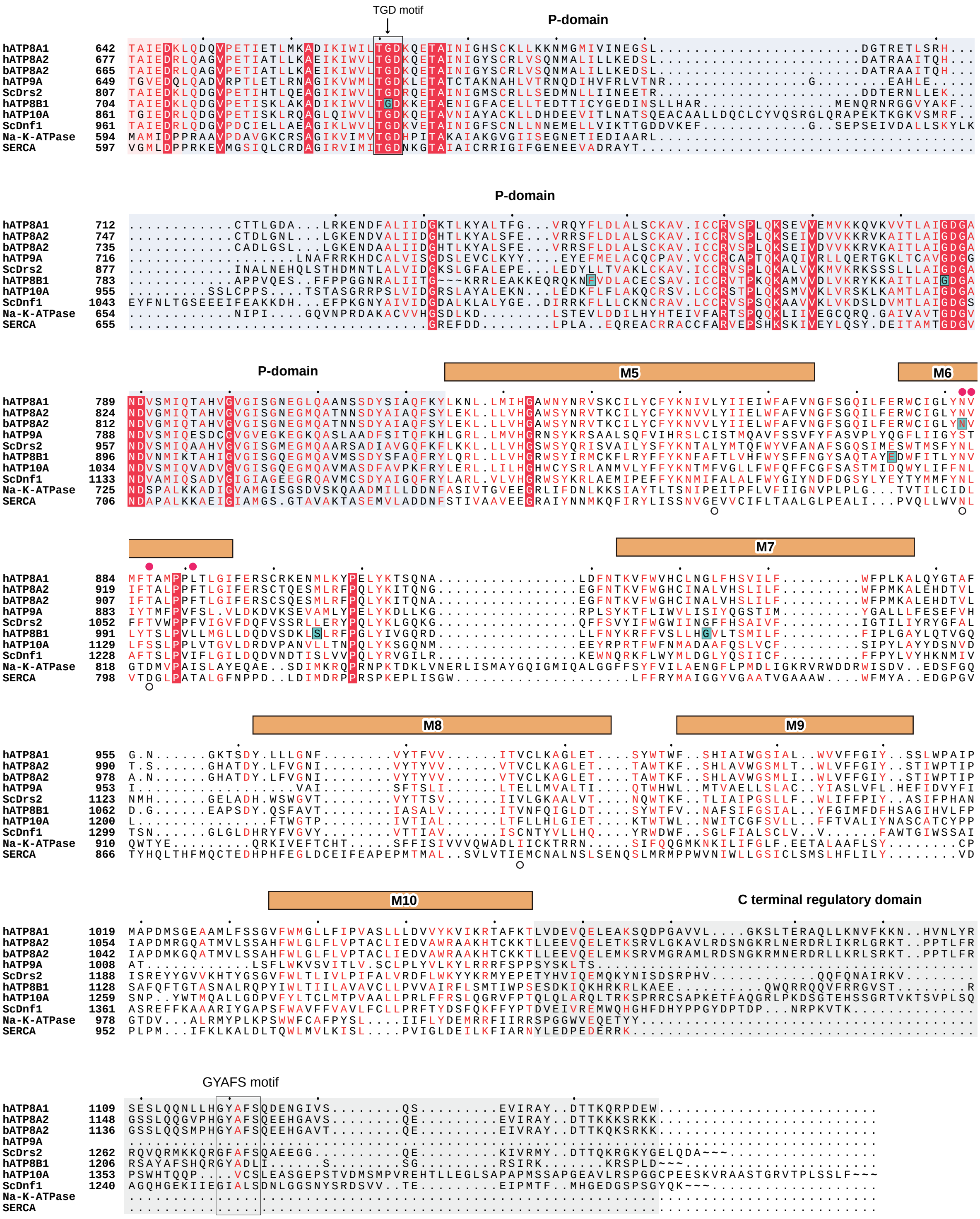


Supplementary Figure 2

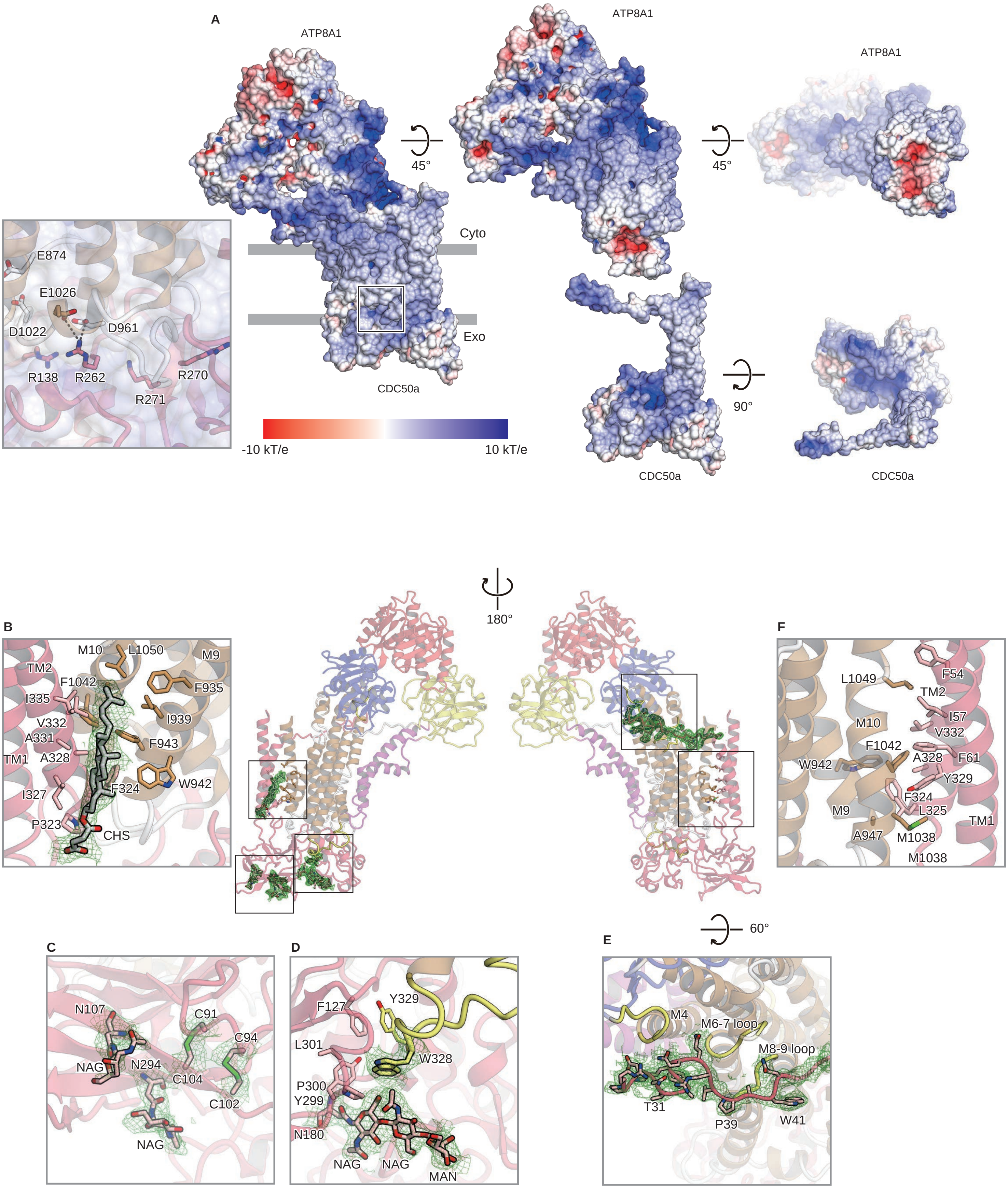


### Supplementary Figure 3

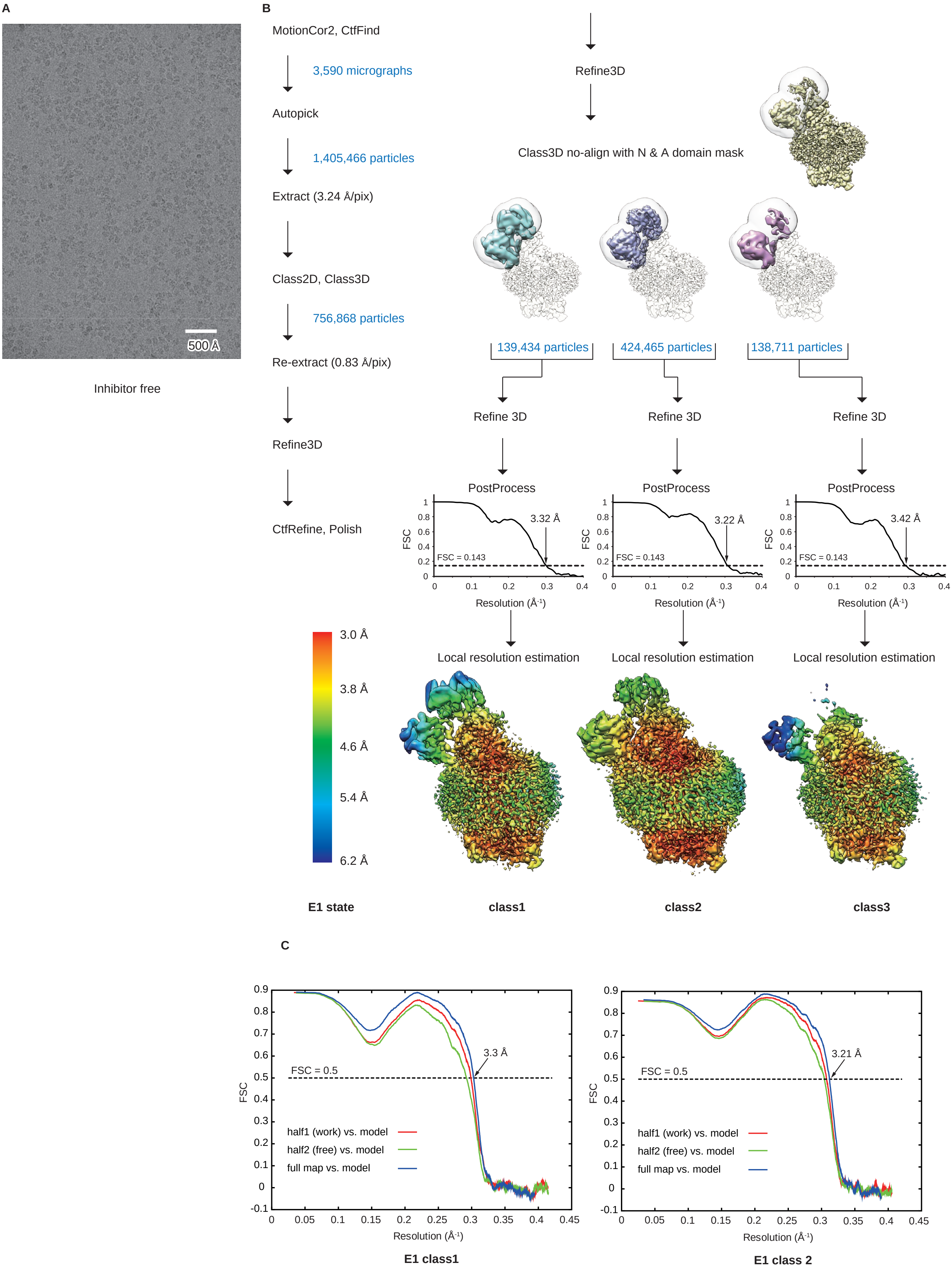


### Supplementary Figure 4

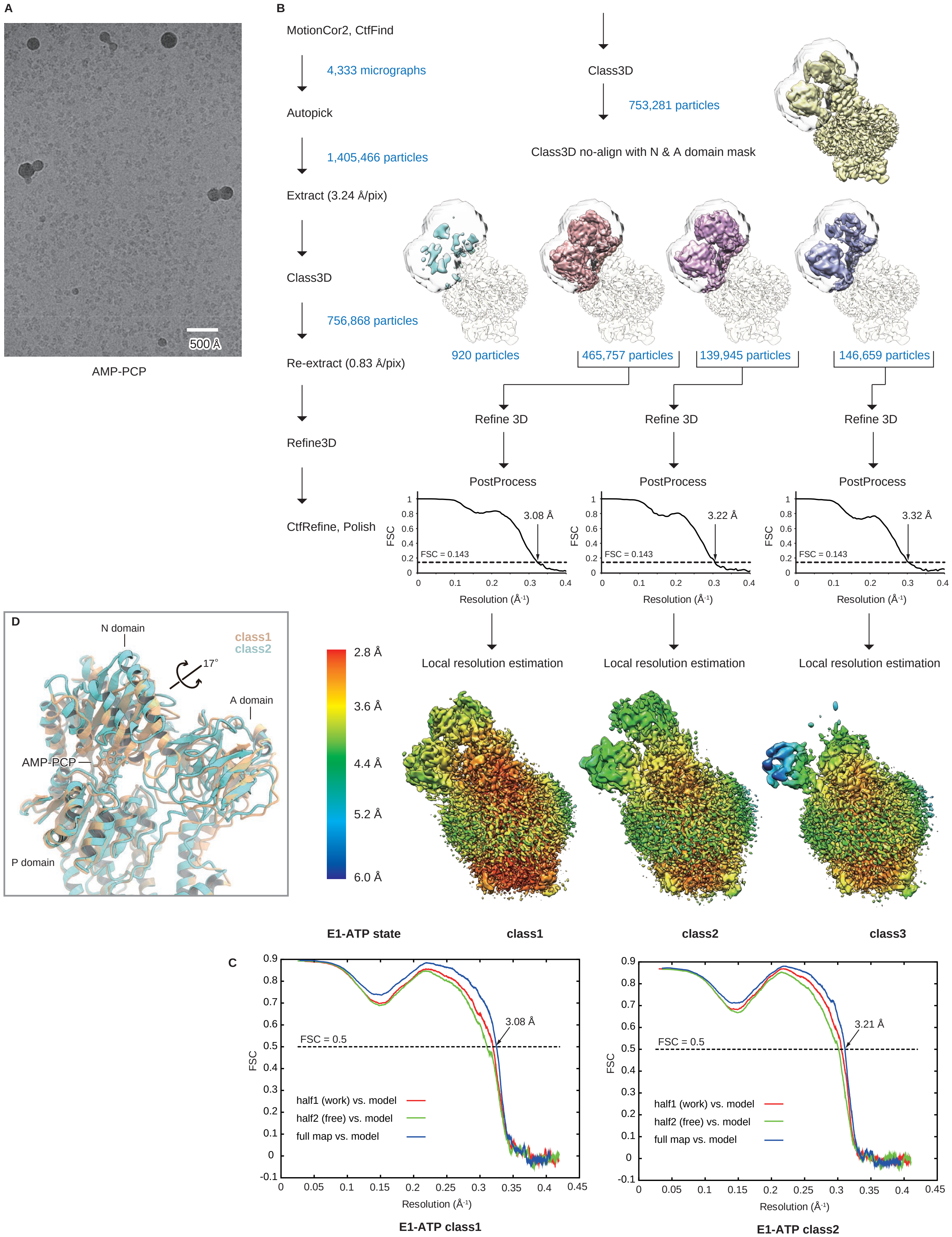


Supplementary Figure 5

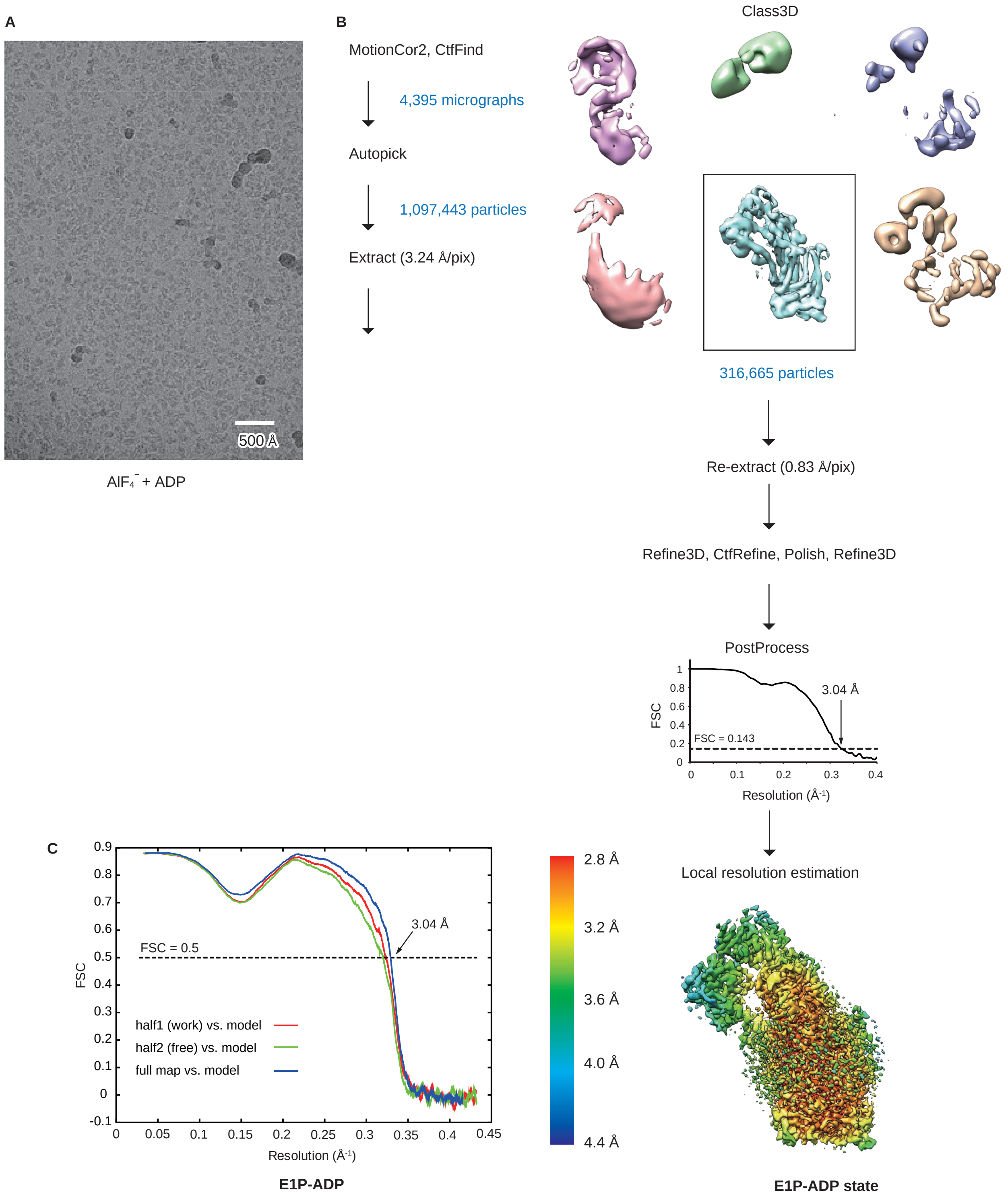


### Supplementary Figure 6

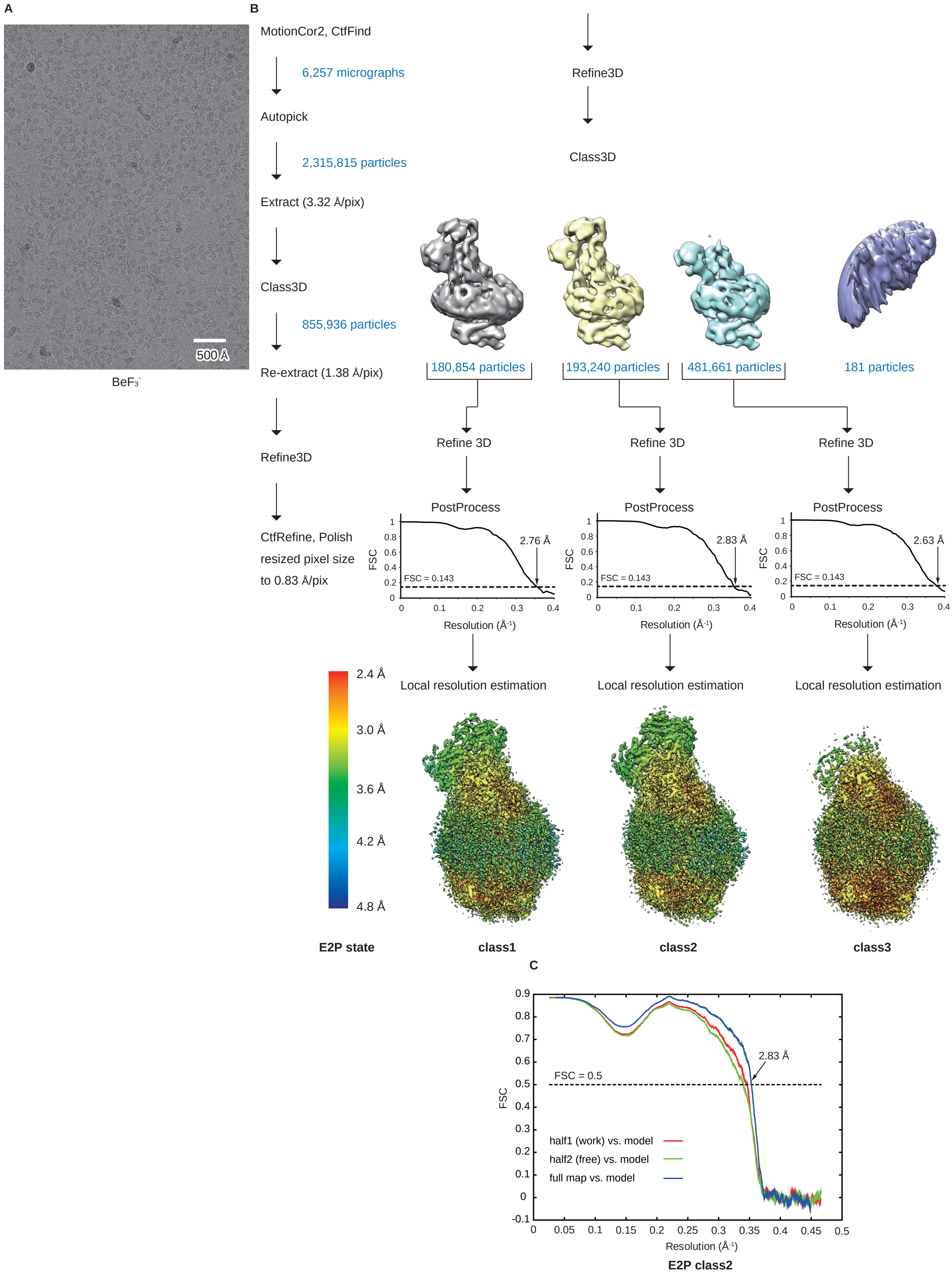


### Supplementary Figure 7

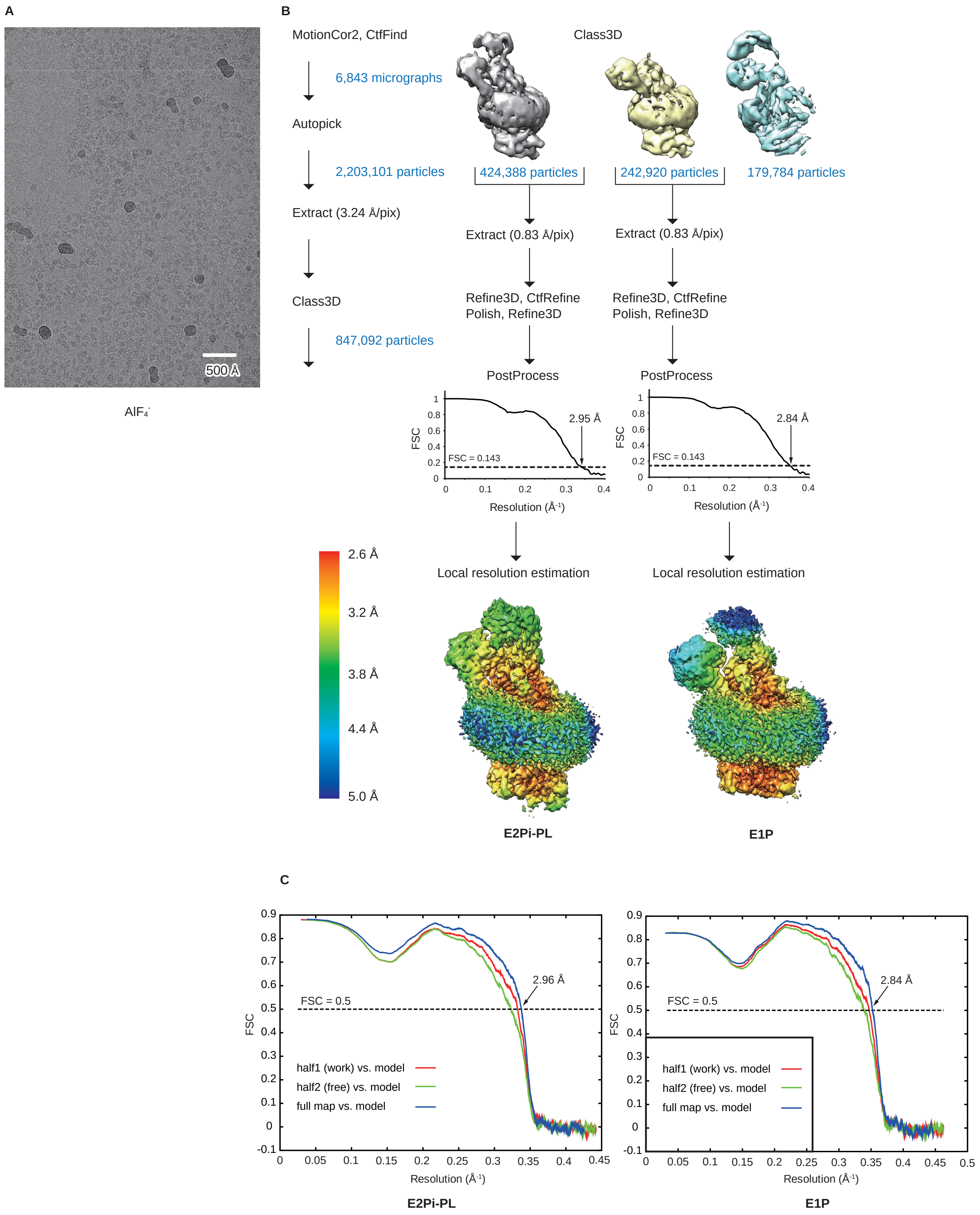


Supplementary Figure 8

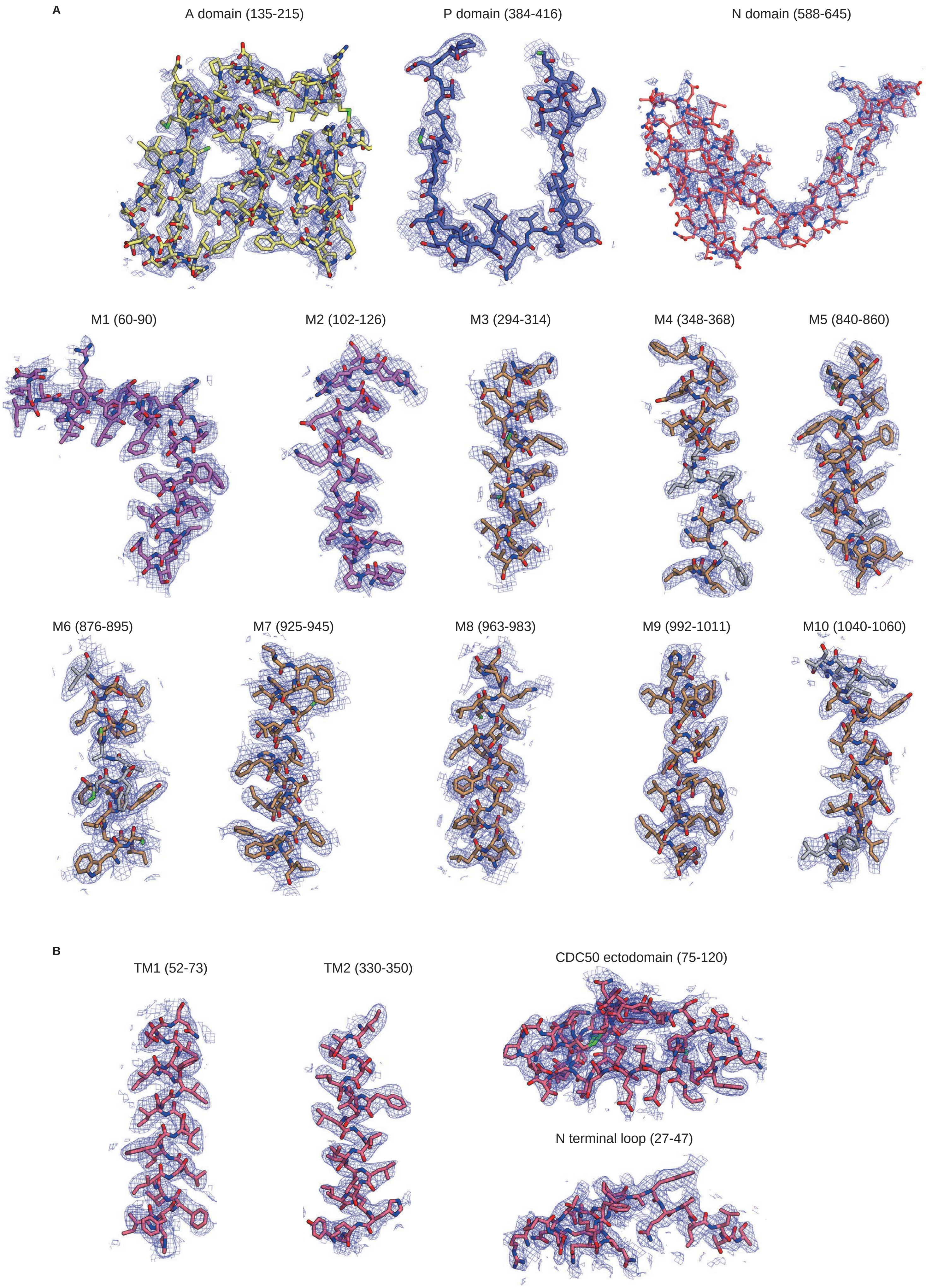


Supplementary Figure 9

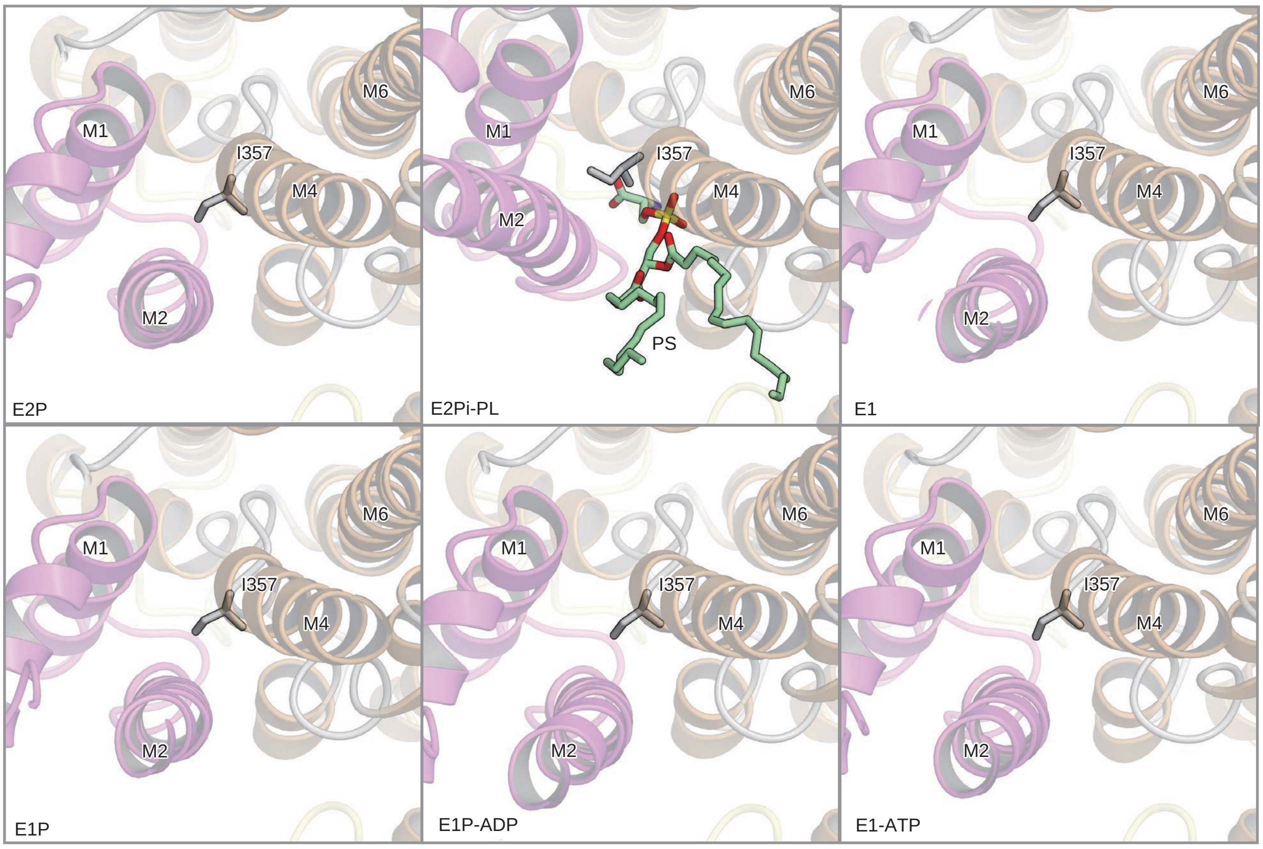


### Supplementary Figure 10

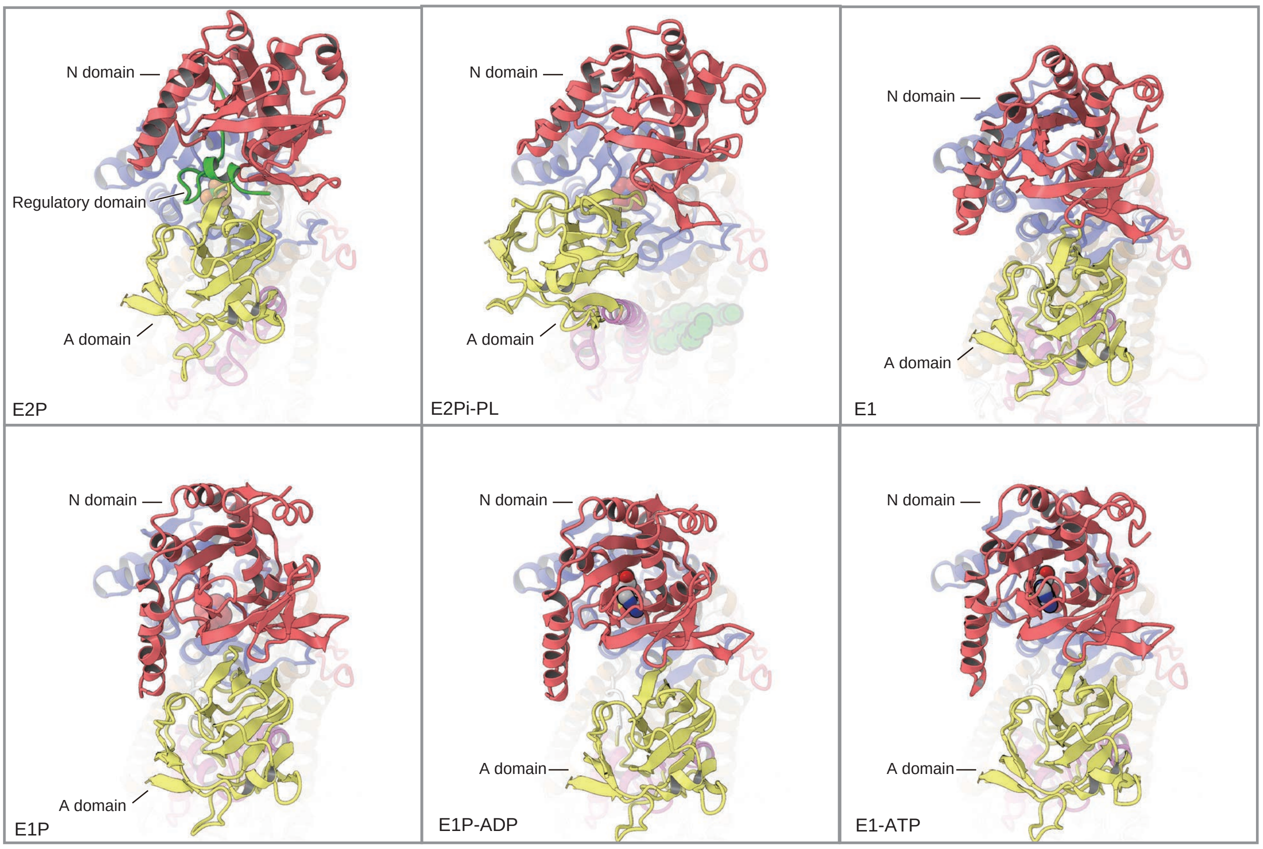


### Supplementary Figure 11

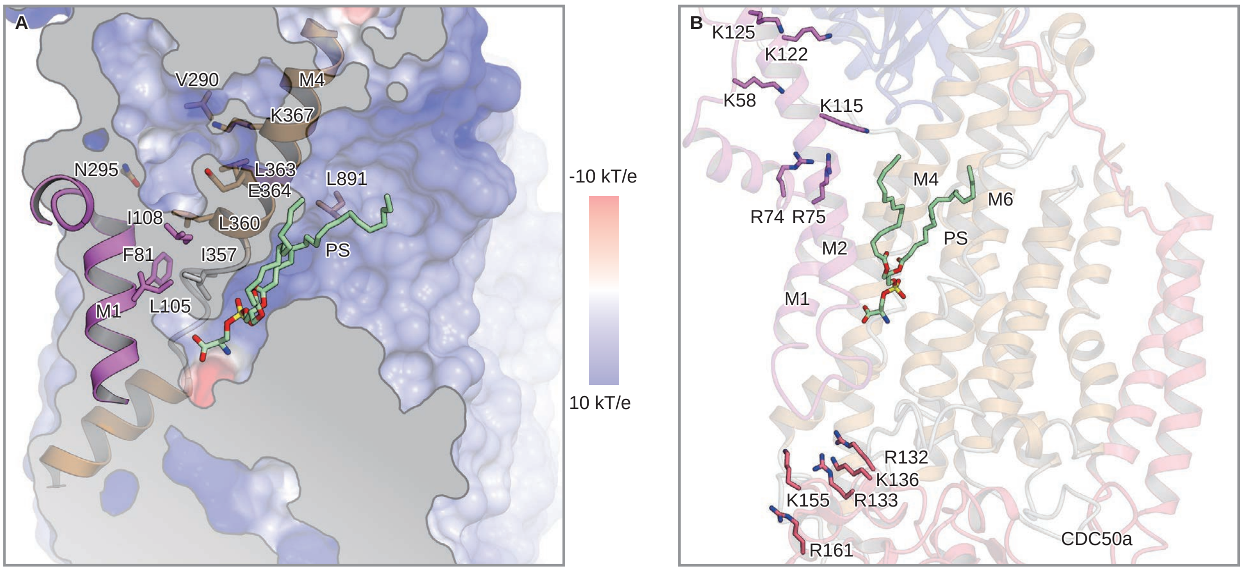


### Supplementary Figure 12

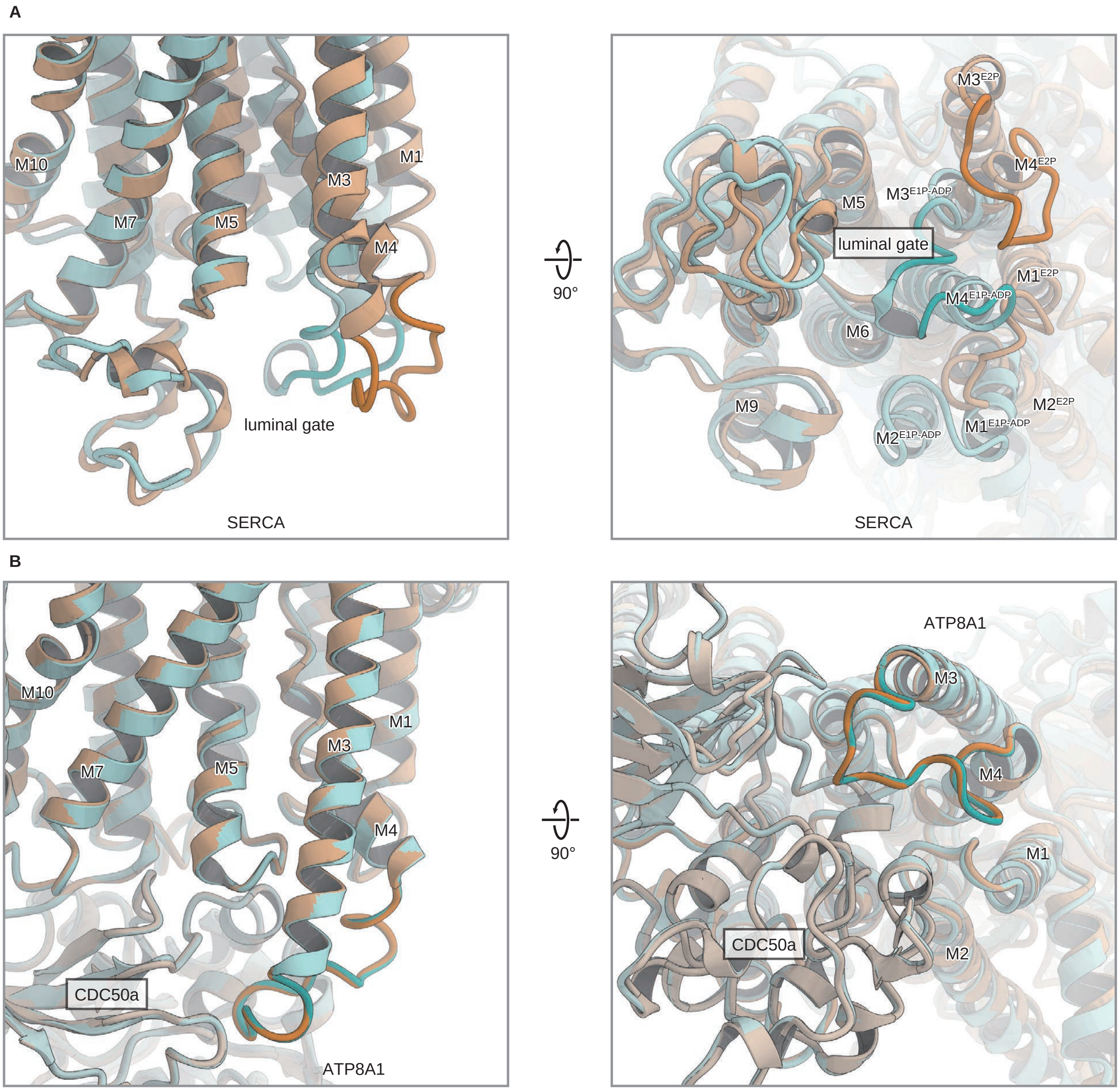


### Supplementary Figure 13

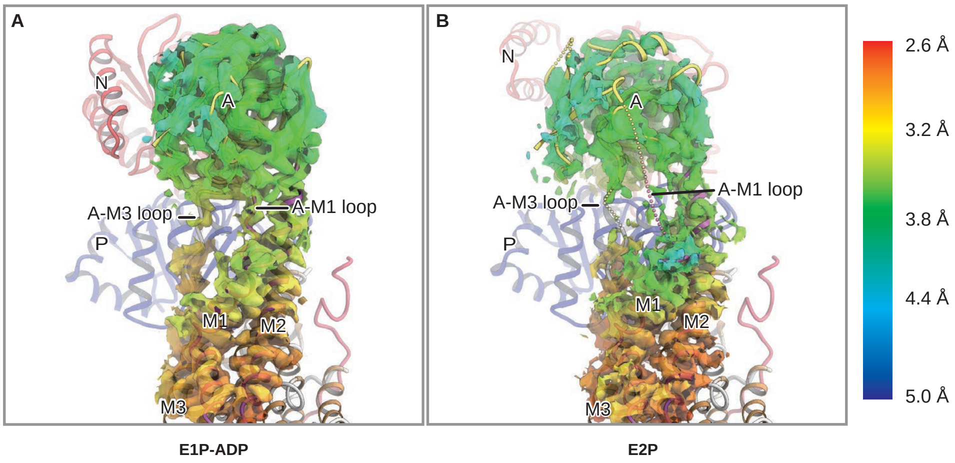


### Supplementary Figure 14

### Cryo-EM data collection, refinement and validation statistics

| **Data collection and processing** | | | | | | | | | | | | |
|---|---|---|---|---|---|---|---|---|---|---|---|---|
| State | E1 class 1 | E1 class 2 | E1 class 3 | E1-ATP class 1 | E1-ATP class 2 | E1-ATP class 3 | E1P-ADP | E2P class 1 | E2P class 2 | E2P class 3 | E2Pi-PL | E1P |
| EMDB-ID | EMD-9931 | EMD-9932 | EMD-9933 | EMD-9935 | EMD-9934 | EMD-9936 | EMD-9937 | EMD-9938 | EMD-9939 | EMD-9940 | EMD-9941 | EMD-9942 |
| PDB ID | 6K7G | 6K7H | | 6K7J | 6K7I | | 6K7K | | 6K7L | | 6K7M | 6K7N |
| Inhibitor | No | | | AMP-PCP | | | ALF$_4^-$-ADP | BeF$_3^-$ | | | ALF$_4^-$ | |
| Microscope | Titan Krios G3i | | | | | | | | | | | |
| Detector | K3 Summit | | | | | | | | | | | |
| Magnification | 105,000 | | | | | | | | | | | |
| Voltage (kV) | 300 | | | | | | | | | | | |
| Electron exposure (e$^-$/Å$^2$) | 64 | | | | | | | | | | | |
| Defocus range (μm) | -0.8 to -1.6 | | | | | | | | | | | |
| Pixel size (Å) | 0.83 | | | | | | | | | | | |
| Symmetry imposed | C1 | | | | | | | | | | | |
| Number of movies | 3,590 | | | 4,333 | | | 4,395 | 6,257 | | | 6,843 | |
| Initial particle images | 1,417,719 | | | 1,405,466 | | | 1,097,443 | 2,315,815 | | | 2,203,101 | |
| Final particle images | 139,434 | 424,465 | 138,711 | 465,757 | 139,945 | 146,659 | 316,665 | 180,854 | 193,240 | 481,661 | 424,388 | 242,920 |
| Map resolution (Å) | 3.32 | 3.22 | 3.42 | 3.08 | 3.22 | 3.32 | 3.04 | 2.76 | 2.83 | 2.63 | 2.95 | 2.84 |
| FSC threshold | 0.143 | 0.143 | 0.143 | 0.143 | 0.143 | 0.143 | 0.143 | 0.143 | 0.143 | 0.143 | 0.143 | 0.143 |
| Map resolution range (Å) | 3.05 to 6.17 | 2.94 to 5.67 | 3.14 to 6.96 | 2.81 to 5.52 | 2.95 to 5.65 | 3.01 to 5.80 | 2.76 to 5.62 | 2.42 to 4.89 | 2.58 to 5.29 | 2.55 to 5.99 | 2.67 to 5.29 | 2.60 to 5.30 |
| Map sharpening B factor (Å$^2$) | -94.5 | -92.0 | -99.3 | -112.0 | -99.1 | -102.3 | -88.6 | -59.4 | -64.9 | -71.4 | -71.2 | -84.3 |
| **Model building and refinement** | | | | | | | | | | | | |
| Model resolution (Å) | 3.3 | 3.21 | | 3.08 | 3.21 | | 3.04 | | 2.83 | | 2.96 | 2.84 |
| FSC threshold | 0.5 | 0.5 | | 0.5 | 0.5 | | 0.5 | | 0.5 | | 0.5 | 0.5 |
| Model composition | | | | | | | | | | | | |
| Protein atoms | 10,421 | 10,421 | | 10,416 | 10,416 | | 10,412 | | 10,446 | | 10,379 | 10,396 |
| Metals | 0 | 0 | | 1 | 0 | | 3 | | 2 | | 2 | 2 |
| Other atoms | 101 | 101 | | 132 | 132 | | 132 | | 104 | | 157 | 105 |
| R.m.s. deviations | | | | | | | | | | | | |
| Bond lengths (Å) | 0.012 | 0.014 | | 0.012 | 0.011 | | 0.012 | | 0.006 | | 0.011 | 0.015 |
| Bond angles (degree) | 1.200 | 1.226 | | 1.244 | 1.210 | | 1.218 | | 1.034 | | 1.177 | 1.373 |
| **Validation** | | | | | | | | | | | | |
| Clashscore | 5.88 | 5.55 | | 5.02 | 5.06 | | 5.82 | | 4.83 | | 5.46 | 6.65 |
| Rotamer outliers (%) | 0.7 | 0.5 | | 0.3 | 1.1 | | 0.8 | | 0.5 | | 0.4 | 0.5 |
| Ramachandran plot | | | | | | | | | | | | |
| Favored (%) | 88.9 | 88.8 | | 88.4 | 88 | | 89.2 | | 90.5 | | 88.5 | 88.8 |
| Allowed (%) | 10.8 | 11 | | 11.4 | 11.8 | | 10.7 | | 9.2 | | 11.2 | 10.8 |
| Outlier (%) | 0.3 | 0.2 | | 0.2 | 0.2 | | 0.2 | | 0.2 | | 0.3 | 0.4 |

### Supplementary Data Table 1
